# Nuclear Translocation of Vitellogenin in the Honey Bee (*Apis mellifera*)

**DOI:** 10.1101/2021.08.18.456851

**Authors:** Heli Salmela, Gyan Harwood, Daniel Münch, Christine Elsik, Elías Herrero-Galán, Maria K. Vartiainen, Gro Amdam

**Author notes:** Corresponding author: Gyan Harwood, Department of Entomology, University of Illinois at Urbana-Champaign, 320 Morrill Hall, 505 South Goodwin Avenue, Urbana, IL 61801.

## Abstract

Vitellogenin (Vg) is a conserved protein used by nearly all oviparous animals to produce eggs. It is also pleiotropic and performs functions in oxidative stress resistance, immunity, and, in honey bees, behavioral development of the worker caste. It has remained enigmatic how Vg affects multiple traits. Here, we asked whether Vg enters the nucleus and acts via DNA-binding. We used immunohistology, cell fractionation and cell culturation to show that a structural subunit of honey bee Vg translocates into cell nuclei. We then demonstrated Vg-DNA binding theoretically and empirically with prediction software and chromatin immunoprecipitation with sequencing (ChIP-seq), finding binding sites at genes influencing immunity and behavior. Finally, we investigated the immunological and enzymatic conditions affecting Vg cleavage and nuclear translocation, and constructed a 3D structural model. Our data are the first to show Vg in the nucleus and suggests a new fundamental regulatory role for this ubiquitous protein.

## 1. Introduction

Vitellogenin (Vg) is an egg yolk precursor protein common to nearly all oviparous animals and likely evolved around 700 million years ago as the oldest member of the Large Lipid Transfer Protein (LLTP) family (Hayward et al., 2010). It is highly pleiotropic, functioning as a storage protein (Amdam and Omholt, 2002), an immunomodulator (Du et al., 2017; Garcia et al., 2010; Li et al., 2008), an antioxidant (Havukainen et al., 2013; Salmela et al., 2016), and a behavior regulator (Amdam et al., 2006; Antonio et al., 2008; Nelson et al., 2007). These functions are observed in taxa as disparate as fish, corals, and insects, suggesting that Vg may have evolved to support multiple functions. Honey bees (*Apis mellifera*) have been the premier insect model for studying Vg pleiotropy, owing largely to the central role it plays in colony division of labor. Here, the non-reproductive worker caste performs a series of age-dependent tasks that is regulated, in part, by Vg titers circulating in their hemolymph (Nelson et al., 2007).

The cause of Vg’s pleiotropy remains somewhat enigmatic. We have detailed understanding on how Vg forms egg yolk (Tufail and Takeda, 2008), and how it acts in innate immunity as a pathogen pattern recognition receptor (Li et al., 2008), but the molecular mechanism(s) by which it influences multiple traits remains unclear. Some pleiotropic proteins implement their multiple effects by acting as transcription factors via DNA-binding or participating in gene-regulatory complexes that affect the expression of many genes (Chesmore et al., 2016). Interestingly, down-regulation of honey bee Vg by means of RNA-interference mediated *vg* gene knockdown alters the expression of thousands of genes (Wheeler et al., 2013). However, no previous research has addressed or established whether Vg can translocate into the cell nucleus and bind DNA. This lack of investigation may be due to Vg’s large size, as the proteins are typically around 200 kDa (Tufail and Takeda, 2008), while most passively-transported nuclear proteins are below 60 kDa (Wang and Brattain, 2007). Yet in many animals, Vg is enzymatically cleaved and reassembled prior to secretion (Tufail and Takeda, 2008). In invertebrates such as insects, Vg is cleaved in the vicinity of a multiply phosphorylated polyserine linker sequence stretch that resides between two evolutionarily conserved Vg protein domains called the N-sheet and the α-helical domain (Baker, 1988; Havukainen et al., 2012; Tufail and Takeda, 2008). In honey bees, this cleavage occurs *in vivo* in the fat body, the primary production site of Vg (Tufail and Takeda, 2008), and results in a detached 40 kDa N-sheet (Havukainen et al., 2011). The function of this N-sheet is not fully known, but in fish it serves as the receptor binding region (Li et al., 2003).

In this study, we asked whether pleiotropic effects of Vg may be partly explained by nuclear translocation and DNA-binding of the conserved N-sheet, and we used the honey bee as our study organism. First, we used confocal microscopy and Western blots of nuclear compartments to test whether the N-sheet subunit can translocate into the nucleus of fat body cells, using an antibody targeting the honey bee N-sheet. We verified these results by observing nuclear uptake of fluorescently labelled Vg in cell culture. Second, we predicted the DNA-binding capability of honey bee Vg using sequence- and structure-based software, and then used ChIP-seq to identify Vg binding sites in honey bee DNA. We compared workers <24 h old and 7 days old to determine whether Vg-DNA binding sites persist as workers transition between behavioral tasks. We searched for *de novo* binding motifs and performed a gene ontology analysis of Vg-DNA binding sites, identifying many sites associated with immune- and behavior-related genes. These data motivated a functional response-to-challenge assay using *E. coli* to reveal if behavior of Vg cleavage and translocation is, in fact, dynamic. Finally, we established the enzymatic conditions required for Vg cleavage, and we developed a three-dimensional structure model to elucidate the critical N-terminal area of the protein.

## 2. Material and Methods

### 2.1 Antibody against honey bee Vg 40 kDa N-sheet

The honey bee Vg 40 kDa fragment (Uniprot ID Q868N5; the amino acid residues 24-360) was produced in *E. coli* by GenScript (Piscataway, NJ, USA); it was subcloned into pUC57 vector, and an N-terminal hexahistidine tag was used for one-step affinity purification. The polyclonal serum antibody was raised in a rat by Harlan Laboratories (Boxmeer, the Netherlands), and tested by Western blotting against proteins extracted from the honey bee (see S1).

### 2.2 Immunohistology

N=6 winter worker bees were collected from two hives at Norwegian University of Life Sciences, Aas, Norway. The gut was removed, and the abdomen detached and placed for overnight fixing in 4% paraformaldehyde in 4 ºC, followed by three PBS washes, 10 min each. The fat body was dissected in PBS and de/rehydrated with a full ethanol series. For each individual, one tissue sample was used as test (primary and secondary antibodies) and another one as control (no primary antibody). The samples were incubated with the primary antibody overnight in 4°C (1:50 in 2 % BSA-PBS-Triton X-100). DAPI was used as a nuclear marker. The anti-rat secondary antibody was Alexa-568 nm that has no emission spectrum overlap with DAPI. For qualitative anatomical descriptions, a high NA (=high resolution) objective was chosen (40x immersion oil; NA 1.25). Scans were taken with zoom 1.0 and zoom 2.0. All images were taken in sequential acquisition mode to minimize crosstalk between the two channels for detecting DAPI and the Vg signal.

### 2.3 Cell fractionation

Nine winter worker honey bee individuals were anesthetized in cold and the fat body tissue was dissected in ice cold PBS. Tissues were placed in three tubes with 50 µl hypotonic buffer (20 mM Tris-HCl pH 7, 10 mM NaCl and 3 mM MgCl_2_) pooling three individuals per tube. 2.5 µl 10 % NP-40 was added, followed by 10 s vortex and centrifugation for 10 min 3000 g in 4°C. The supernatant was collected as the cytosolic fraction. The pellet was washed once with 500 µl hypotonic buffer, and then suspended in 30 µl hypotonic buffer with 5 mM EDTA, 1 % Tween-20 and 0.5 % SDS and vortexed. All samples were then centrifuged for 10 min 15000 g in 4°C. 15 µl of each sample was run on SDS-PAGE gel and blotted.

### 2.4 Western blotting

The gel and Western blot reagents were purchased from Bio-Rad, and the protocol for all blots was as follows: The blotted nitrocellulose membrane was incubated with PBS containing 0.5% Tween with 2.5% bovine serum albumin overnight. The membrane was incubated for 1 h with the N-sheet antibody (1:2000) and for 1 h with a horseradish peroxidase-conjugated secondary antibody (1:5000) prior to imaging using an Immun-Star kit. All gels and blots were imaged, and band intensities were measured using a ChemiDoc XRS imager (Bio-Rad).

### 2.5 Whole Vg purification

Vg was purified from wintertime worker honey bee hemolymph as in Salmela *et al*. 2015.

### 2.6 Labeled Vg in cell culture

Pure Vg was labeled with Alexa Fluor 488 protein labeling kit (Invitrogen, Carlsbad, CA, USA). HighFive cells were grown on an 8-well chamber slide (Thermo Fisher) overnight (number of slides one, repeated three times). The media was replaced with 20 µg/µl Vg-488 in PBS and incubated in dark for 1 h. The cells were washed twice with PBS, fixed with 4 % paraformaldehyde and the washes were repeated. DAPI was used as a nuclear marker. The cells were imaged the following day with Leica TCS SP5 MP (63x).

### 2.7 DNA-binding prediction

The following prediction tools available for protein-DNA binding were tested with default settings: Sequence-based DNABIND (Liu and Hu, 2013), DP-BIND (Hwang et al., 2007) and DRNApred (Yan and Kurgan, 2017), and structure-based DNABIND (Liu and Hu, 2013) and DISPLAR (Tjong and Zhou, 2007). The structure used was the honey bee Vg N-sheet homology model published earlier (Havukainen et al., 2011).

### 2.8 ChIP-seq

To test empirically whether Vg binds to DNA, and to determine what types of genes and genomic regions Vg is bound to, we performed ChIP-seq on newly emerged (<24 hours) and 7-day old worker bees. Newly emerged workers will take on tasks such as cleaning comb cells, while workers at 7 days old act as nurses to nourish the queen and developing larvae. Thus, workers in these two age groups span an important behavioral transition or maturation that is integral to a colony’s division of labor. Samples were created by pooling fat body tissue from 10 individuals from each age group, all originating from the same hive. Freshly harvested samples were flash frozen in liquid nitrogen and homogenized. ChIP was carried out with established protocols (Bai et al., 2013) using Dynabeads™ Protein G (Invitrogen). We opted to use polyclonal antibodies raised in rabbits against the whole 180 kDa Vg molecule (Pacific Immunology, Ramona, CA) (Jensen and Børgesen, 2000) rather than the rat-origin antibodies against the 40 kDa Vg N-terminal domain used elsewhere in this study because of their superior immunoprecipitation performance. In a preliminary study, the rabbit-origin whole-Vg antibodies consistently pulled down more chromatin than multiple batches of the rat-origin Vg N-terminal antibodies, which failed to retrieve sufficient chromatin for sequencing. This is likely due to Protein G (and A) having a greater affinity for rabbit-origin than rat-origin antibodies, as per the manufacturer. As a negative control, we compared immunoprecipitated DNA against input DNA (i.e., DNA from the same sample not precipitated with antibodies). We opted against using anti-GFP antibodies in our negative control, as this type of negative control is less commonly used and more prone to background noise (Xu et al., 2021).

Chromatin samples were sequenced at the DNASU lab at Arizona State University. The raw Illumina 2×75bp pair-end reads were quality checked using FastQC v0.10.1 (Andrews, 2010), followed by adapter trimming and quality clipping by Trimmomatic 0.35 (Bolger et al., 2014). Any reads with start, end or the average quality within 4bp window falling below quality scores 18 were trimmed. The clean reads were aligned to reference genome *Apis mellifera* Amel_4.5 (https://www.ncbi.nlm.nih.gov/genome/48?genome_assembly_id=22683) by Bowtie 2 version 2.2.9 (Langmead and Salzberg, 2012). Another round of QC was performed on bam files. Library complexity was checked by NRF (non-redundancy fraction), defined as the number of unique start positions of uniquely mappable reads divided by number of uniquely mappable reads. All the samples passed the threshold 0.8 recommended by ENCODE. IGVtools and bamCompare from deepTools (Ramírez et al., 2014) were employed to compare two BAM files based on the number of mapped reads. First the genome is partitioned into bins of equal size and then the number of reads in each bin is counted. The log2 value for the ratio of number of reads per bin was reported for IGV visualization. MACS2 was used for peaks calling with 0.01 FDR cutoff. Narrowpeak files as MACS2 output were annotated by HOMER (Heinz et al., 2010). It first determines the distance to the nearest TSS and assigns the peak to that gene. Then it determines the genomic annotation of the region covered by the center of the peak, including TSS, TTS, Exon, Intronic, or Intergenic.

To test for non-random occurrence of peaks within genome features, we used 1000 random peak datasets from the 7-day old worker data set. To create the random peak datasets, we used the shuffle program of the BEDTools package (Aronesty, 2011) on the 782 peaks from the 7-day data set that were located on full chromosome assemblies to generate 1000 bed files with peak locations that were shuffled within chromosomes, such that shuffled peak locations were non-overlapping and did not occur in assembly gaps. The annotatePeaks.pl program from the Homer package (Heinz et al., 2010) was then used to annotate the 1000 shuffled peak data sets and the 7-day peak data set with respect to genome features in the NCBI *Apis mellifera* RefSeq annotation. We used chi-squared tests to determine whether the observed numbers of peaks overlapping promoter plus Transcription Start Site (TSS) regions (−1 kb to +100 bp from TSS), Transcription Termination Sites (TTS), exons, introns and intergenic regions were greater or less than expected by chance.

### 2.9 Gene ontology

We performed a gene ontology (GO) term analysis to determine if our list of Vg-DNA binding sites was enriched for any biological processes, molecular functions, cellular components, or KEGG pathways. We used the list of sites from 7-day old bees, but omitted sites found in *intergenic* regions. Honey bee genes were converted to *D. melanogaster* orthologues (N = 360 genes), as this latter genome is more robustly annotated. We used DAVID 6.8 (Huang et al., 2009) with default settings and the resulting terms are those deemed significant (P < 0.05) after Benjamini-Hochberg corrections.

### 2.10 Vg-DNA binding motif search

For *de novo* motif identification, we created a non-redundant dataset of 790 peaks by combining peaks identified in the newly emerged and 7-day old samples and merging peaks with overlapping regions between the two datasets. We used GimmeMotifs (Heeringen et al., 2011) which ran and combined results for ten motif prediction packages – Mdmodule, Weeder, GADEM, trawler, Improbizer, BioProspector, Posmo, ChIPMunk, JASPAR, AMD, HMS and Homer. GimmeMotifs clusters the results, performs enrichment and computes receiver operator characteristic (ROC) and mean normalized conditional probability (MNCP). Half of the peaks (395) were used to train each algorithm and the other half was used for testing. Since we did not have *a priori* knowledge of the motif length, we ran GimmeMotifs four times with different size range options (small 5-8 bp, medium 5-12 bp, large 6-15 bp, xl 6-20 bp). Since the peaks were located throughout the genome, we used sequences randomly chosen from the genome with GC content to the peak sequences as the background for the enrichment tests.

### 2.11 Vg response to immune challenge, enzymatic cleavage, and 3D structural model

See S2.

## 3. Results

### 3.1. Vg translocation to the nucleus

#### 3.1.1. Immunohistology

We observed fat body cells using confocal microscopy (Fig 1A-E). Honey bee fat body contains two cell types: Trophocytes are characterized by irregularly-shaped nuclei and are responsible for Vg production and storage, while oenocytes have spherical nuclei and do not produce or contain Vg (Pan et al., 1969; Roma et al., 2010). We observed two Vg localization patterns in trophocytes: i) Vg signal co-localized with DAPI in the nucleus, and also found spread throughout the cytosol (Fig 1A-B), and ii) Vg signal absent in the nucleus and instead restricted to granule-like formations in the cytosol (Fig 1C, see also (Havukainen et al., 2011) for observations of this pattern). Controls for autofluorescence and unspecific secondary antibody staining were included (Fig 1D-E), which indicated that the immunosignals present in Fig 1A-D are specific for Vg antibody incubation.

**Fig 1.**
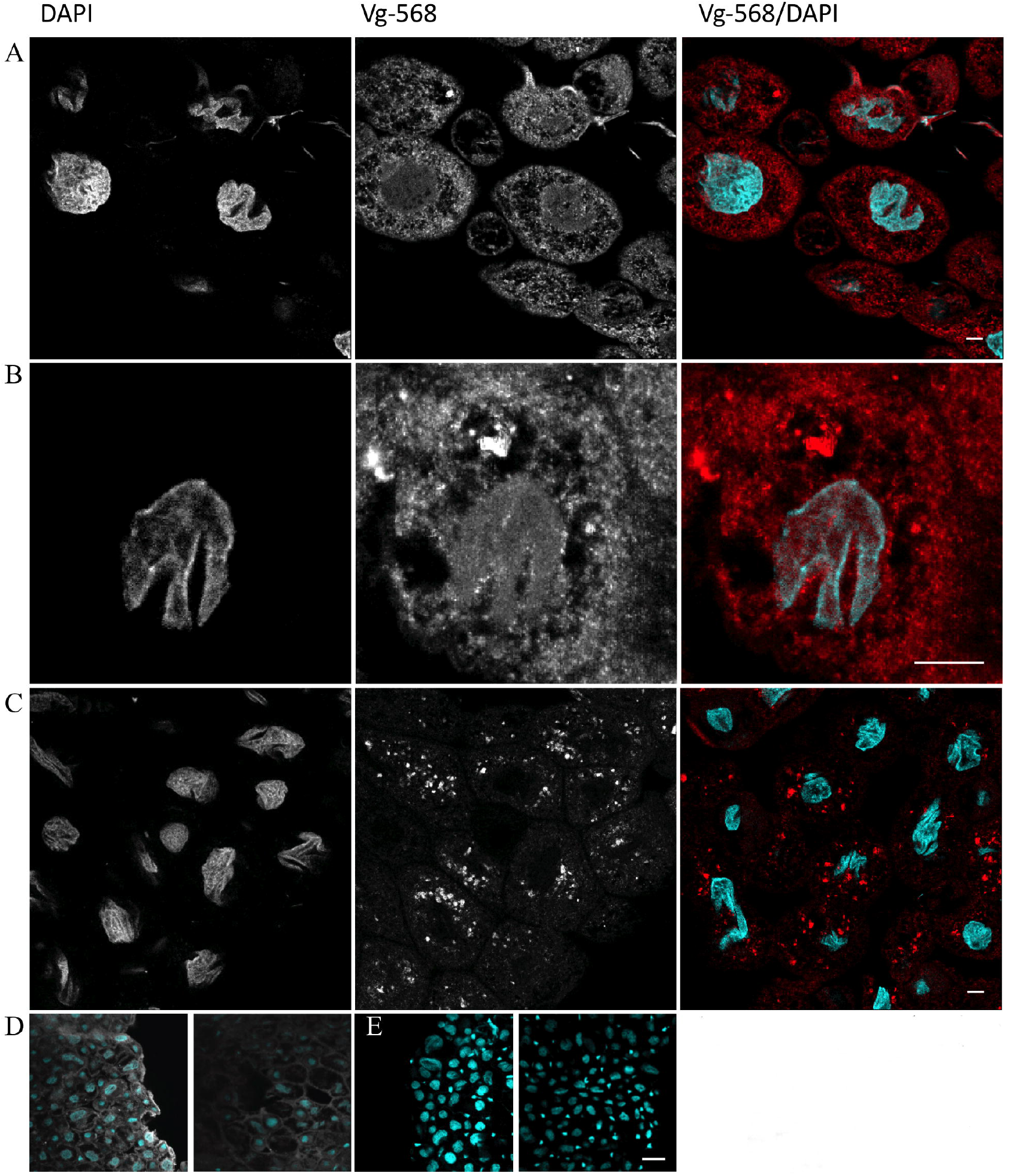
Localization of Vg signals in the honey bee fat body using an antibody targeting the N-terminal 40 kDa domain. A-E Confocal images of fat body cells representing six samples. The last panel shows the superposition of DAPI (cyan) and Vg signal (red). A: Vg signal co-localizes with DAPI in cell nuclei. B: Zoom-in of a single cell showing Vg nuclear translocation. C: Cells that do not show Vg co-localization with DAPI, instead, Vg signal is found in granules in the cytosol. The scale bar = 10 µm. D-E show representative images for controls where the primary antibody was applied (D) or omitted (E). Note that when omitting Vg antibody incubation, no immunosignal was detected (grayscale), while only the DAPI signal (cyan) was present. Scalebar = 50 µm. Magnification 40x.

#### 3.1.2. Cell fractionation

To verify the nuclear location of the Vg N-sheet, we separated fat body cells into cytosolic and nuclear compartments by cell fractionation followed by western blot. Full-length Vg (180 kDa) and a ∼75 kDa band were found in the cytosolic component, while the nuclear component only contained fragments below 75 kDa, including the 40 kDa N-sheet domain (Fig 2). We have shown previously that the 40 kDa Vg subunit is a specific cleavage product of fat body cells and not simply a biproduct of unspecific degradation (Havukainen et al., 2012, 2011). We also observed 70, 37 and 25 kDa fragments in the nuclear fraction, but these are likely degradation products caused by the method, as the bands are faint or non-existent in untreated Western blot samples (see S1).

**Fig 2.**
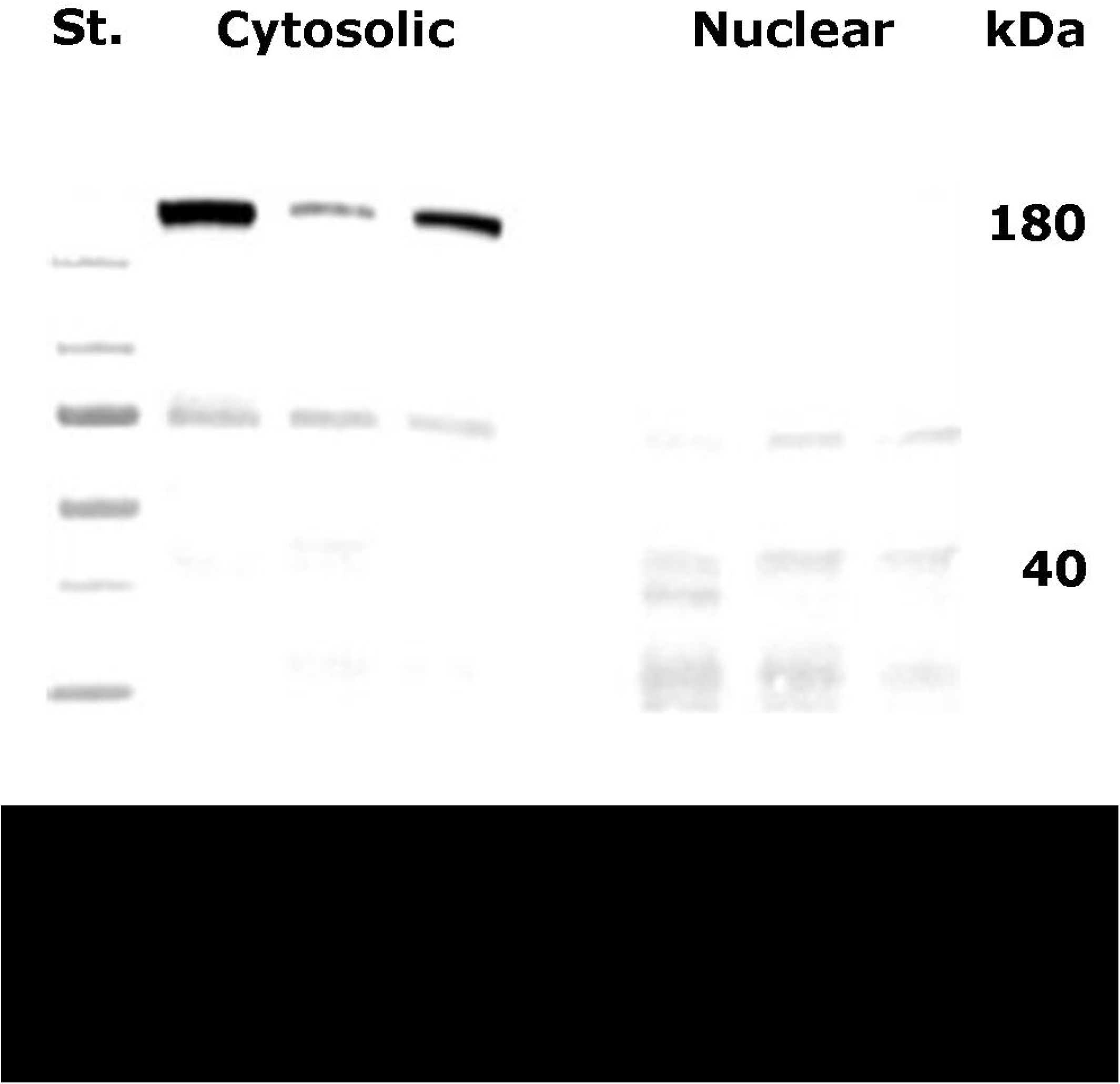
Western blot of cytosolic and nuclear fractions of honey bee fat body tissue. St. = size standard. The full-length (180 kDa) Vg is dominating the cytosolic fraction of the fat body proteins, whereas the 40-kDa N-terminal fragment mostly localizes in the nuclear fraction. Also, other fragments of approximately 70 and 25 kDa were visible in the nuclear fraction. Three individuals were pooled for each lane. N = 3.

#### 3.1.3. Labeled whole Vg in cell culture

To rule out that the nuclear signal was an artefact caused by Vg primary antibody, we incubated cell cultures (Lepidopteran ovarian HighFive cells) with antibody-free pure fluorescently-labelled Vg. During incubation, the fluorescent Vg was imported into cells and was visible in the cell nuclei co-localizing with DAPI, and also visible in the cytosol in granule-like formations (Fig 3).

**Fig 3.**
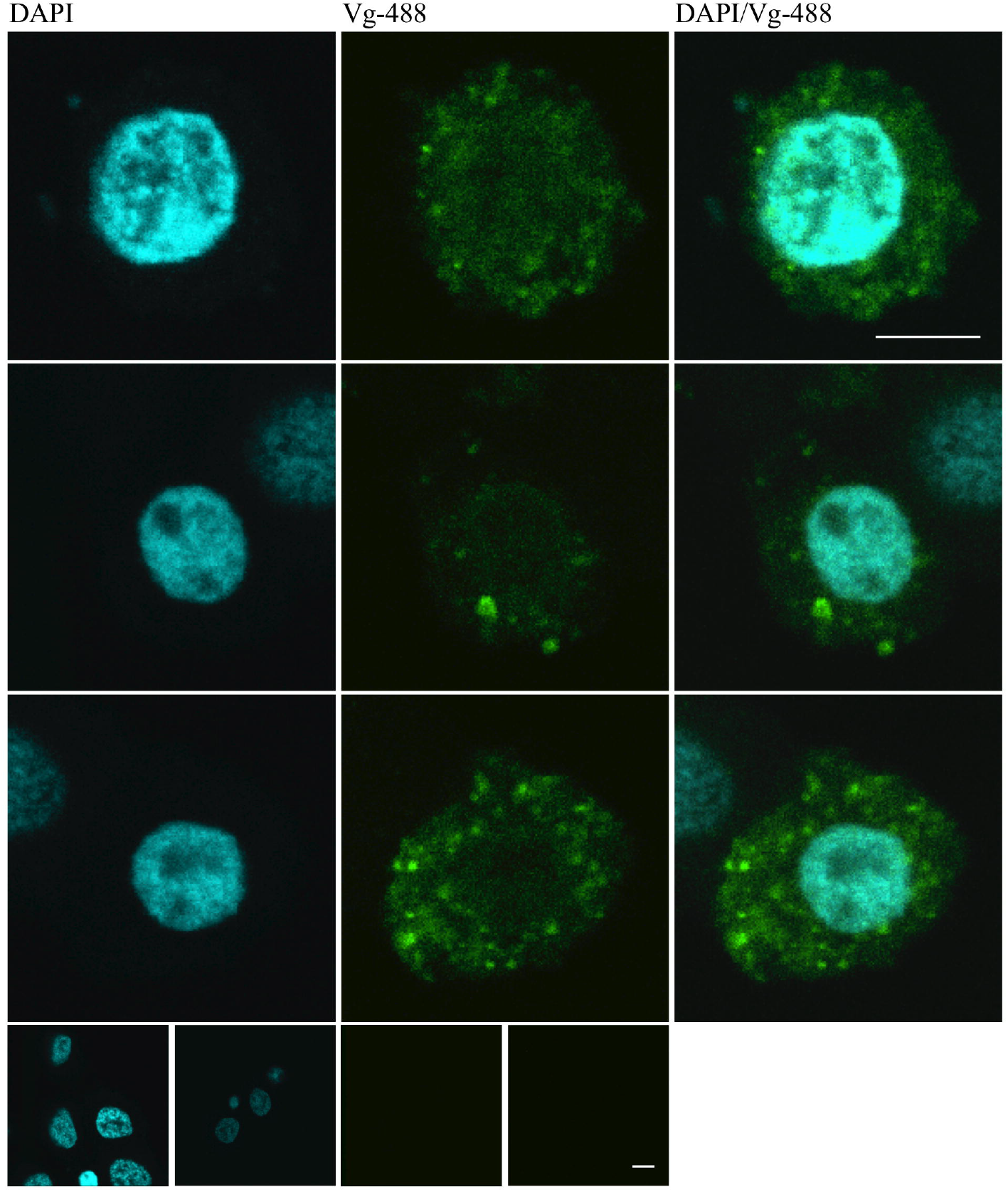
Localization of Alexa-488-labelled purified Vg in cultured insect HighFive cells. Vg was observed both in cytosol mostly in bright granules and as haze in the nucleus in the confocal sections. The lowest row shows two Vg-negative controls imaged with the same settings. Scale bar = 5 µm. Magnification 63x.

### 3.2 Vg-DNA binding

#### 3.2.1. In silico binding prediction

Using the whole Vg amino acid sequence, two separate programs with different search algorithms, DP-Bind (Hwang et al., 2007) and DRNApred (Yan and Kurgan, 2017), both identified the same amino acid residue stretch as a putative DNA binding domain: SRSSTSR in position 250-256 of the N-sheet domain of Vg. Another sequence-based program, DNABIND (Liu and Hu, 2013), also identified N-terminal Vg as a DNA-binding protein. This program predicts a protein’s DNA binding probability and sets a threshold probably of 0.5896, above which a protein is statistically likely to be able to bind DNA. Vg scored a 0.6267 probability, indicating it can bind DNA. Additionally, the structure-based prediction software DISPLAR (Tjong and Zhou, 2007) identified the SRSSTSR stretch as capable of binding DNA using a published honey bee Vg N-sheet model (Havukainen et al., 2011) as input. There were another 12 and 3 additional stretches identified by DP-BIND and DRNApred, respectively, which did not overlap between the programs, whereas DISPLAR identified two additional stretches that did not overlap with predictions made by the two sequence-based programs. Taken together, results from multiple prediction software platforms support the hypothesis that Vg can bind to DNA, and this capability is likely restricted to the N-sheet.

#### 3.2.2 ChIP-seq

ChIP DNA had significant enrichment (FDR < 0.01) at 90 and 782 putative Vg-DNA binding sites on fully assembled chromosomes in newly emerged and 7-day old workers, respectively. Of the 90 binding sites in newly emerged bees, 83 (92%) were also present in 7-day old bees, illustrating an expansion of Vg-DNA binding sites as workers age. Using the 7-day old worker binding sites, we analyzed their genomic distribution compared to a null distribution of random peaks and found a greater number than expected by chance within promotor-TSS, TTS, and intergenic regions, and fewer than expected by chance in exons and introns (Fig 4).

**Fig 4.**
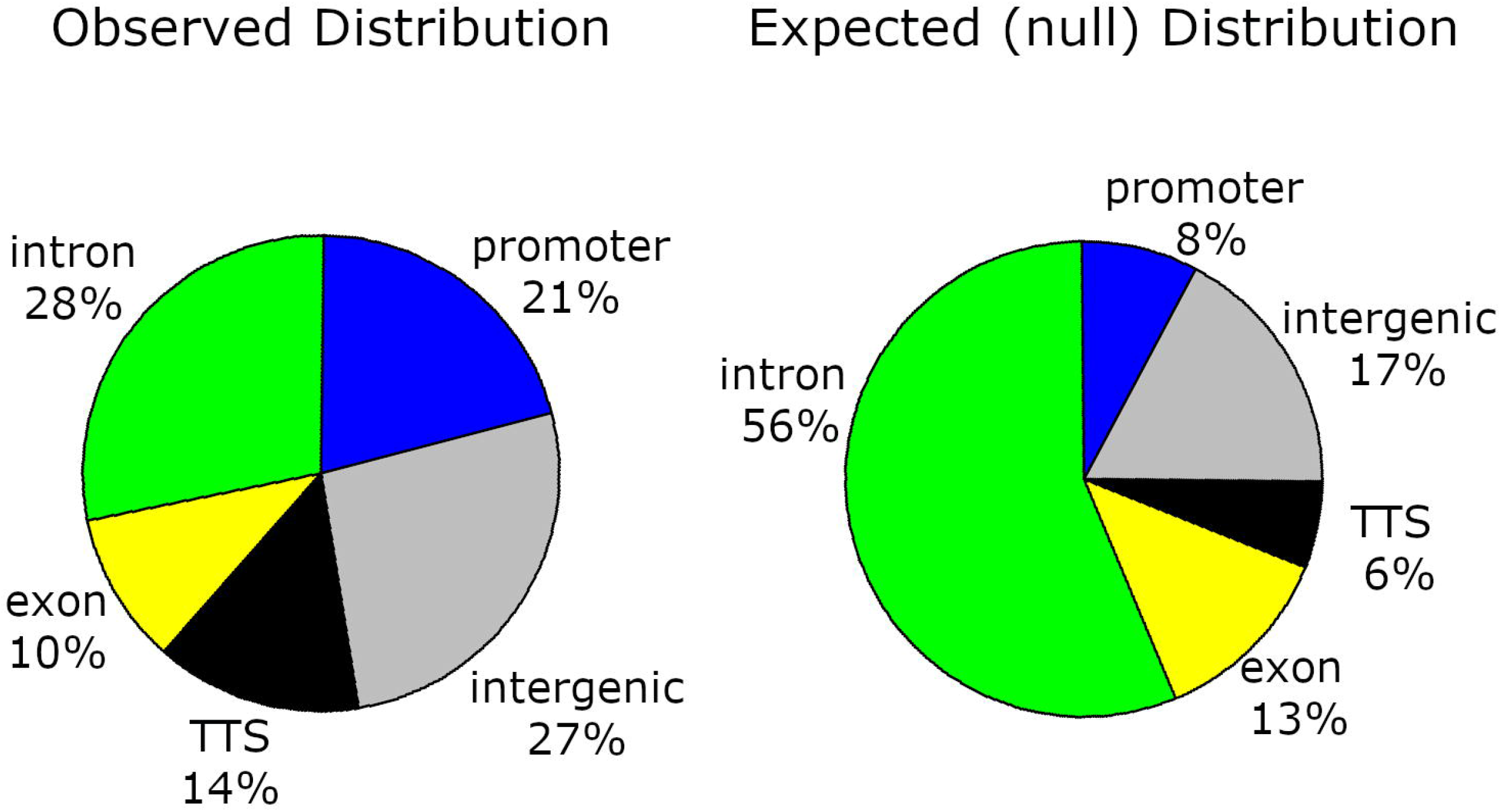
Genomic distribution of observed Vg-DNA binding sites compared with a null distribution. The null distribution was created by randomly shuffling 782 non-overlapping ChIP-seq peaks for 1000 iterations. We found more Vg-DNA binding sites than expected by chance in promotor regions (χ^2^ = 175.18, P = 5.47E-40), transcription termination sites (χ^2^ = 91.68, P = 2.09E-24), and intergenic regions (χ^2^ = 46.14, P = 1.10E-11), and fewer than expected by chance in exons (χ^2^ = 3.75, P = 0.05) and introns (χ^2^ = 244.55, P = 4.01E-55)

#### 3.2.3. Gene Ontology

Next, we again used the 7-day old worker binding sites to perform a Gene Ontology (GO)-term analysis. We omitted binding sites located in intergenic regions and restricted our list to sites located in or near genes (promotor regions, introns, exons, tts, N=574 binding sites across 484 genes). We converted these *Apis mellifera* genes into *Drosophila melanogaster* orthologs (N=274 genes) using the Hymenoptera Genome Database (Elsik et al., 2016) and used DAVID Bioinformatics Resources (Huang et al., 2009) to look for enrichment of biological processes, molecular functions, cellular components, and KEGG pathways. We found significant enrichment (Benjamini-Hochberg-corrected P value < 0.05) for 142 terms and pathways (see Table S1). There was much overlap in these terms, but most genes had functions pertaining to *developmental processes* (GO:0032502, N=134 genes, adj. P = 6.58 E-8), *neurogenesis* (GO:0022008, N=70 genes, adj. P = 4.63 E-8), *signaling* (GO:0023052, N=75 genes, adj. P = 4.74 E-5), or similar functions. Many genes bound by Vg code for membrane-bound receptors that are known to play roles in complex phenotypes, like behavior, including receptors for corazonin, glutamate, acetylcholine, serotonin, and octopamine (see Table S1 for complete list of Vg binding sites). Furthermore, Vg was bound to more than a dozen immunological genes, including *Defensin-1*, a key antimicrobial peptide, and *sickie*, an integral component of the Immune Deficiency pathway defense against gram-negative bacteria. These results demonstrate that Vg binds to DNA at hundreds of loci in the honey bee genome, that many are behaviorally-, developmentally-, and immunologically-relevant, and that many of these Vg-DNA interactions persist as adult workers transition between tasks.

#### 3.2.4 Binding motif identification

We analyzed the binding site data to look for DNA sequences to which Vg may be binding. We ran four analyses using GimmeMotif (Heeringen et al., 2011), and we selected the top motif predictions, based on a combination of percent enrichment, P-value, ROC-AUC and MNCP (Table 1) (Fig 5). The top motif prediction for the small motif run was a sub-motif of the large motif run, so it was discarded. These results suggest that there are specific sequence motifs to which Vg binds.

**Table 1.**
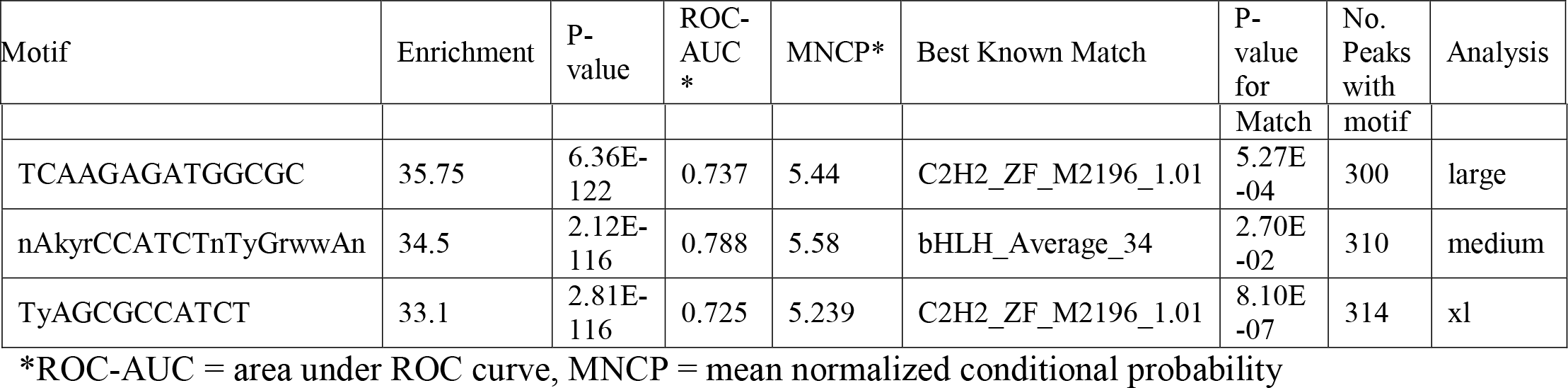
Top de novo motif predictions.

**Fig 5.**
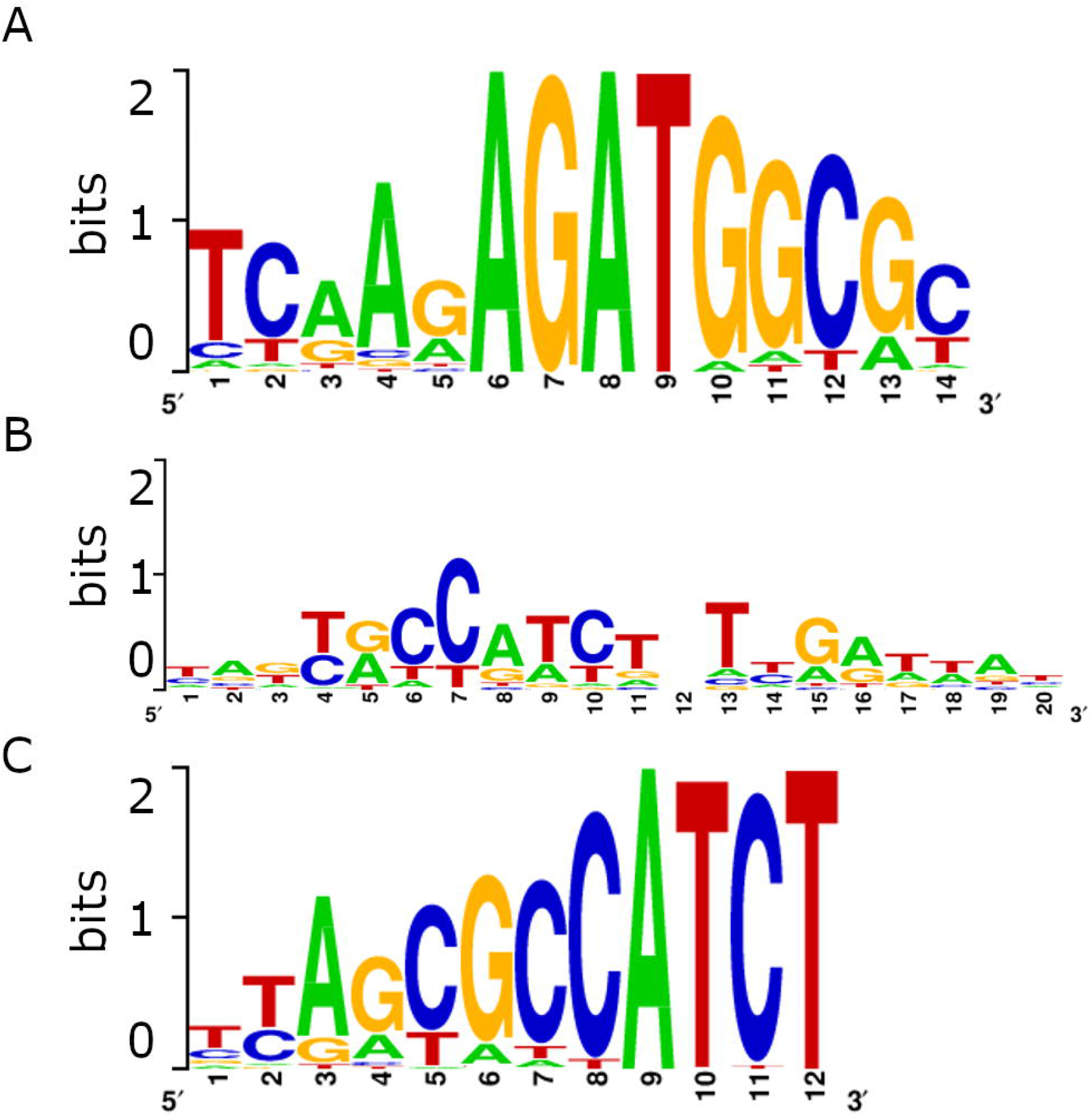
Logo representations for the top motif predictions. A) Top motif prediction from analysis with “large” motif setting (35.75% enrichment). B) Top motif prediction from analysis with “medium” motif setting (34.5% enrichment). C) Top motif prediction from analysis with “xl” motif setting (33.1% enrichment).

### 3.3 Vg cleavage dynamics

#### 3.3.1. Vg protein cleavage and nuclear translocation in response to immune challenge

We found that Vg incubated *in vitro* with *E. coli* had increased protein fragmentation, but that worker bees that ingested *E. coli* possessed fewer fat body nuclei with Vg signal within than workers fed a control diet (see S2).

#### 3.3.2. Vg structure and enzymatic cleavage

We found that the linker region connecting the N-sheet with the rest of the Vg molecule protrudes from the structure, providing easy access to proteolytic enzymes. Moreover, enzymatic cleavage is likely carried out, at least partially, via dephosphorylation and caspase activity (see S2).

## 4. Discussion

Vg dates back at least 700 million years to the time when animals first appeared (Hayward et al., 2010) (Fig 6), and the ubiquity of its roles in reproduction and immunity across diverse taxa (Du et al., 2017; Li et al., 2008; Salmela et al., 2015) suggest that these functions developed early in its evolutionary history. Several of its structural domains are highly conserved, and yet the molecular mechanism by which Vg regulates so many complex traits like behavior and longevity has remained something of a mystery. Here, we reveal a major and hitherto unknown ability of Vg that may provide such a molecular explanation: nuclear translocation and DNA binding. Vg may be directly involved in modulating gene expression, and given its prevalence in the animal kingdom and its conserved molecular structure, it is possible that it performs a similar function across a wide array of oviparous animals.

**Fig 6.**
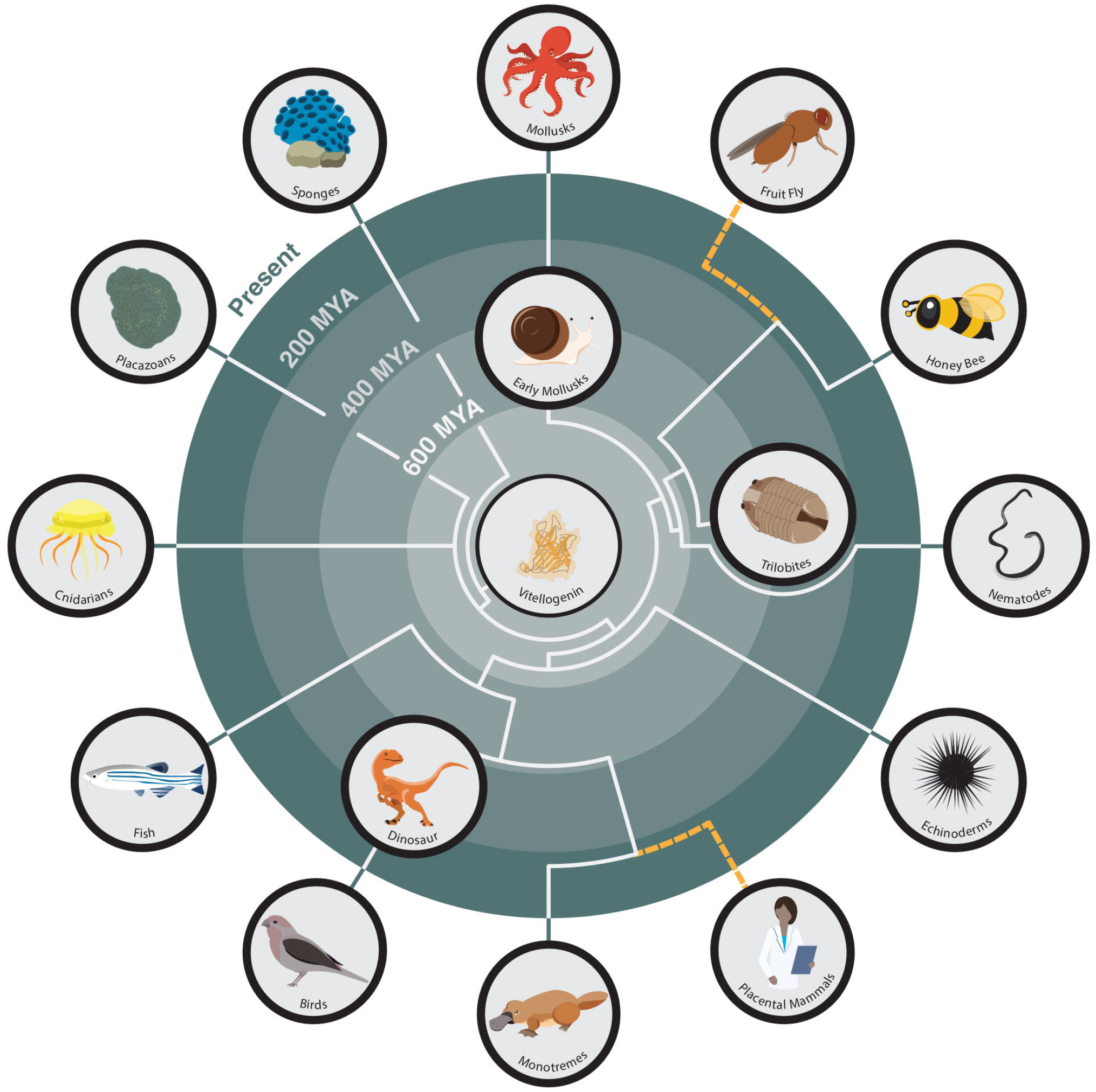
The deep phylogenetic history of Vitellogenin. Vg first evolved around 700 million years ago when Metazoans appeared (Hayward et al., 2010) and is present in all extant Metazoan phyla, from earliest animals like sponges and cnidarians to the more recently evolved chordates, like fish, birds, and monotreme mammals (Agnese et al., 2013; Akasaka et al., 2013; Babin, 2008; Biscotti et al., 2018; Chen et al., 2018, 1997; García-Alonso et al., 2006; Prowse and Byrne, 2012; Riesgo et al., 2014). Vg has been lost in several scientifically important lineages, including placental mammals and higher dipterans like *D. melanogaster* (depicted with dashed yellow lines) (Brawand et al., 2008; Sappington, 2002). Vg’s earliest known functions pertain to egg-yolk formation and immunity, but it remains to be seen when DNA binding evolved.

In our study, we found strong evidence of Vg translocating to the nucleus using three methods: immunohistology, Western blot of proteins in the fractionated nuclear compartment, and antibody-free localization of Vg in cell culture. The two first approaches confirmed the Vg N-sheet domain to be present in both nucleus and cytosol, with the latter result reaffirming previous findings (Havukainen et al., 2011). Moreover, the second approach specifies that the cell nucleus excludes the full length 180 kDa Vg molecule and the previously reported specific Vg fragmentation product of 150 kDa size (i.e., the Vg molecule without the N-sheet domain) (Havukainen et al., 2011). The cell culture approach makes it less likely that antibody artefacts explain the outcomes of the first experiments.

We found theoretical support for Vg’s DNA binding ability using several prediction software platforms, and this prompted us to test this ability empirically. Our ChIP-seq analysis confirmed that honey bee Vg binds to many loci in fat body DNA of newly emerged and 7-day old workers. Despite there being far fewer Vg-DNA binding sites in newly emerged than 7-day old workers (150 vs 927 loci, respectively), there is robust overlap in binding sites between these two age groups as over 90% of sites observed in newly emerged workers are still present in 7-day older workers. This suggests that Vg not only maintains long-term associations with specific loci, but also that the repertoire of Vg-DNA binding sites expands as workers age and behaviorally transition from cell cleaning to nursing.

GO term analysis of Vg-DNA binding sites hint at the types of biological functions that Vg may be targeting for regulation. We found significant enrichment for the GO terms *developmental processes* (GO:0033502), *neurogenesis* (GO:0022008), and *signaling* (GO:0023052), among many others, and many of the genes included within these terms code for membrane proteins. Indeed, many genes code for G protein-coupled receptors and ion channels (Table S1), suggesting that Vg regulates components of signal transduction pathways. Interestingly, newly emerged and 7-day old workers perform different tasks, and many of these receptors are known to affect how individuals behave and respond to stimuli. These include receptors for glutamate (Kucharski et al., 2007), acetylcholine (Eiri and Nieh, 2012), corazonin (Gospocic et al., 2017), and octopamine (Grohmann et al., 2003). One possible explanation is that as workers age and transition between tasks they respond to different signals from nestmates and the environment, and thus must be equipped with relevant molecular machinery to receive and respond to those signals.

A potential shortcoming here is that these Vg-DNA binding loci are from fat body cells rather than neurons, so the effect on honey bee behavior might appear limited. However, Vg is neither found nor expressed in honey bee neurons (Münch et al., 2015), limiting its likelihood of binding to DNA therein. Fat body, on the other hand, not only produces and stores Vg, but also plays a central role in regulating metabolism and homeostasis. In this capacity, it produces many products that affect a wide range of insect behaviors, including courtship (Lazareva et al., 2007), host-seeking (Klowden et al., 1987), and onset of foraging (Antonio et al., 2008). Moreover, the fat body is a key player in innate immunity (Fehlbaum et al., 1994; Lycett et al., 2006; Morishima et al., 1997), and our data show Vg to be bound to several important immune-related genes in this tissue. These include *defensin-1*, an important antimicrobial peptide (Ganz, 2003), *sickie*, a regulator of immune transcription factors (Foley and O’Farrell, 2004), and several genes involved in autophagy and phagocytosis (Levine and Deretic, 2007; Strand, 2008)(Table S1). Taken together, the fat body’s central role in honey bee biology means that any protein-DNA binding herein could have wide-ranging effects on numerous physiological pathways. This work also revealed several putative *de novo* DNA binding motifs for Vg. The aim of this ChIP-seq work here was to determine whether Vg binds to DNA, and if so, at which loci. Our next step is to further our understanding of Vg’s actions inside the nucleus and to determine how it regulates gene expression across different caste types and populations in honey bees. This is a multi-faceted approach that will involve performing ChIP-seq on workers, drones, and queens from multiple colonies to determine how Vg differentially binds to DNA, and performing gene expression analyses such as RNA-seq to elucidate how Vg-DNA binding up- or down-regulates gene expression at various loci. Furthermore, we can distinguish whether Vg is a transcription factor or co-regulator by determining what other proteins it interacts with in the nucleus via co-immunoprecipitation and mass spectrometry (Li et al., 2016). This will greatly enhance our knowledge of Vg’s regulatory properties and can potentially unveil co-evolutionary relationships between Vg and other proteins. Nevertheless, the findings presented in this study here represent a major new discovery of Vg function.

Our discovery that the Vg N-sheet subunit binds to DNA in fat body cell nuclei raises several questions that warrant further research. First, does Vg translocate into the nucleus in other tissues in addition to the fat body? Vg has been verified in eggs and ovaries (Seehuus et al., 2007), in hypopharyngeal glands (Seehuus et al., 2007), in immune cells (Hystad et al., 2017), and in glial cells of the honey bee brain (Münch et al., 2015). In this latter observation, it is specifically the Vg N-sheet that is localized in glial cells, and such subcellular localization should prompt a highly relevant research subject since honey bee *vg* knock-downs show an altered brain gene expression pattern (Wheeler et al., 2013) and major behavioral changes (Nelson et al., 2007). Second, does Vg naturally translocate to the nucleus in other animals? The N-terminal Vg cleavage pattern is similar in most insects studied (Tufail and Takeda, 2008), but the possibility of N-sheet translocation remains speculative before it is experimentally tested in another species. Finally, what role do other Vg fragments play in nuclear translocation? The N-sheet-specific antibodies, as well as Vg purified from fat body, show weak protein bands smaller in size to full-length Vg, most notably, bands of ∼75 and ∼125 kDa in size. It is unclear if these are functional Vg fragments or simply the result of unspecific fragmentation. In general, we have observed that Vg fragment number grows in harsh sample treatment conditions. We suspect that at least a 25 kDa fragment detected by the Vg antibody used here in our tissue fractioning assay is a degradation product caused by the assay protocol. However, it is not ruled out that other Vg fragments in addition to N-sheet play a role in nuclear localization of Vg.

Vg was first discovered as an egg-yolk protein, but the protein is so ancient that we cannot be certain of its original function, and it is possible that it assumed a gene regulatory role via DNA-binding early in its history. While we only touch on the mechanistic link between Vg and its multifunctionality, our results will hopefully spark diverse future studies on the role of Vg as a putative transcription factor in honey bees and other animal species.

## Supporting information

Figure S1 - Western blot

Supplementary Material - Additional experimental data

Table S1 - ChIP data and GO term analysis

## 5. Acknowledgements

MSc Taina Stark assisted with the *in vivo* assay. The inhibitors were kind courtesy of Professor Matthew Bogyo at Stanford University, USA. We thank Professor Bogyo and his lab members, as well as Prof. Guy Salvesen (Sanford-Burnham, USA) and Professor Christopher Overall (University of British Columbia, Canada) for advice and innovative discussions. We thank Claus Kreibich (Norwegian University of Life Sciences, Norway), Nicholas Boss (Arizona State University, USA), Osman Kaftanoglu, (Arizona State University, USA), Cahit Ozturk (Arizona State University, USA), Eero Hänninen, Finnish Beekeepers’ Association and Stadin Tarhaajat association for assisting with the live honey bee samples. Thanks to Professor Michelle Krogsgaard (New York University, USA) for help and advice with insect cell culturing, and Dr. Dimitri Stucki for assistance with the statistical test. We also thank Dr. Hua Bai (Iowa State University, USA) and Marc Tatar (Brown University, USA) for insight and training on chromatin immuno-precipitation, as well as Dr. Jason Steele, Dr. Shanshan Yang, and the DNASU Next Generation Sequencing Core (Arizona State University, USA) for sequencing and initial bioinformatics.

## 6. Declarations

### 6.1 Funding

HS was supported by Academy of Finland (#265971). GH was supported by the Natural Sciences and Engineering Research Council of Canada (#376088068) and the School of Life Sciences Research Training Initiative at Arizona State University (#LM5 1079_GH). GA was supported by the Research Council of Norway (#262137).

### 6.2 Conflicts of interest

The authors declare no conflicts of interest.

### 6.3 Availability of data

All relevant experimental data are included in the manuscript and supplementary materials. 3D structural model of Vg is deposited in the Worldwide Protein Data Bank (D_1000249604).

### 6.4 Ethics approval

Not applicable

### 6.5 Consent to participate

Not applicable

### 6.6 Consent for publication

Not applicable

### 6.7 Author’s contributions

GH, HS, and GA conceived this research and designed experiments; HS, GH, DM, CE, EHG and MV performed experiments and analysis; GH and HS wrote the paper; GH made revisions. GA oversaw the project.

## Notes

### Competing Interest Statement

The authors have declared no competing interest.

## References

Agnese, M., Verderame, M., De Meo, E., Prisco, M., Rosati, L., Limatola, E., del Gaudio, R., Aceto, S., Andreuccetti, P., 2013. A network system for vitellogenin synthesis in the mussel Mytilus galloprovincialis (L.). J. Cell. Physiol. 228, 547–555.

Akasaka, M., Kato, K.H., Kitajima, K., Sawada, H., 2013. Identification of Novel Isoforms of Vitellogenin Expressed in Ascidian Eggs. J. Exp. Zoolog. B Mol. Dev. Evol. 320, 118–128. https://doi.org/10.1002/jez.b.22488

Amdam, G.V., Norberg, K., Page Jr., R.E., Erber, J., Scheiner, R., 2006. Downregulation of vitellogenin gene activity increases the gustatory responsiveness of honey bee workers (Apis mellifera). Behav. Brain Res. 169, 201–205. https://doi.org/10.1016/j.bbr.2006.01.006

Amdam, G.V., Omholt, S.W., 2002. The Regulatory Anatomy of Honeybee Lifespan. J. Theor. Biol. 216, 209–228. https://doi.org/10.1006/jtbi.2002.2545

Andrews, S., 2010. FastQC A Quality Control tool for High Throughput Sequence Data [WWW Document]. URL https://www.bioinformatics.babraham.ac.uk/projects/fastqc/ (accessed 10.12.18).

Antonio, D.S.M., Guidugli-Lazzarini, K.R., Nascimento, A.M. do, Simões, Z.L.P., Hartfelder, K., 2008. RNAi-mediated silencing of vitellogenin gene function turns honeybee (Apis mellifera) workers into extremely precocious foragers. Naturwissenschaften 95, 953–961. https://doi.org/10.1007/s00114-008-0413-9

Aronesty, E., 2011. ea-utils: Command-line tools for processing biological sequencing data.

Babin, P.J., 2008. Conservation of a vitellogenin gene cluster in oviparous vertebrates and identification of its traces in the platypus genome. Gene 413, 76–82. https://doi.org/10.1016/j.gene.2008.02.001

Bai, H., Kang, P., Hernandez, A.M., Tatar, M., 2013. Activin Signaling Targeted by Insulin/dFOXO Regulates Aging and Muscle Proteostasis in Drosophila. PLOS Genet. 9, e1003941. https://doi.org/10.1371/journal.pgen.1003941

Baker, M.E., 1988. Is vitellogenin an ancestor of apolipoprotein B-100 of human low-density lipoprotein and human lipoprotein lipase? Biochem. J. 255, 1057–1060.

Biscotti, M.A., Barucca, M., Carducci, F., Canapa, A., 2018. New Perspectives on the Evolutionary History of Vitellogenin Gene Family in Vertebrates. Genome Biol. Evol. 10, 2709–2715. https://doi.org/10.1093/gbe/evy206

Bolger, A.M., Lohse, M., Usadel, B., 2014. Trimmomatic: a flexible trimmer for Illumina sequence data. Bioinformatics 30, 2114–2120. https://doi.org/10.1093/bioinformatics/btu170

Brawand, D., Wahli, W., Kaessmann, H., 2008. Loss of Egg Yolk Genes in Mammals and the Origin of Lactation and Placentation. PLOS Biol. 6, e63. https://doi.org/10.1371/journal.pbio.0060063

Chen, C., Li, H.-W., Ku, W.-L., Lin, C.-J., Chang, C.-F., Wu, G.-C., 2018. Two distinct vitellogenin genes are similar in function and expression in the bigfin reef squid Sepioteuthis lessoniana. Biol. Reprod. 99, 1034–1044. https://doi.org/10.1093/biolre/ioy131

Chen, J.-S., Sappington, T.W., Raikhel, A.S., 1997. Extensive sequence conservation among insect, nematode, and vertebrate vitellogenins reveals ancient common ancestry. J. Mol. Evol. 44, 440–451.

Chesmore, K.N., Bartlett, J., Cheng, C., Williams, S.M., 2016. Complex Patterns of Association between Pleiotropy and Transcription Factor Evolution. Genome Biol. Evol. 8, 3159–3170. https://doi.org/10.1093/gbe/evw228

Du, X., Wang, X., Wang, S., Zhou, Y., Zhang, Y., Zhang, S., 2017. Functional characterization of Vitellogenin_N domain, domain of unknown function 1943, and von Willebrand factor type D domain in vitellogenin of the non-bilaterian coral Euphyllia ancora: Implications for emergence of immune activity of vitellogenin in basal metazoan. Dev. Comp. Immunol. 67, 485–494. https://doi.org/10.1016/j.dci.2016.10.006

Eiri, D.M., Nieh, J.C., 2012. A nicotinic acetylcholine receptor agonist affects honey bee sucrose responsiveness and decreases waggle dancing. J. Exp. Biol. 215, 2022–2029. https://doi.org/10.1242/jeb.068718

Elsik, C.G., Tayal, A., Diesh, C.M., Unni, D.R., Emery, M.L., Nguyen, H.N., Hagen, D.E., 2016. Hymenoptera Genome Database: integrating genome annotations in HymenopteraMine. Nucleic Acids Res. 44, D793–D800. https://doi.org/10.1093/nar/gkv1208

Fehlbaum, P., Bulet, P., Michaut, L., Lagueux, M., Broekaert, W.F., Hetru, C., Hoffmann, J.A., 1994. Insect immunity. Septic injury of Drosophila induces the synthesis of a potent antifungal peptide with sequence homology to plant antifungal peptides. J. Biol. Chem. 269, 33159–33163.

Foley, E., O’Farrell, P.H., 2004. Functional Dissection of an Innate Immune Response by a Genome-Wide RNAi Screen. PLOS Biol. 2, e203. https://doi.org/10.1371/journal.pbio.0020203

Ganz, T., 2003. Defensins: antimicrobial peptides of innate immunity. Nat. Rev. Immunol. 3, 710–720. https://doi.org/10.1038/nri1180

Garcia, J., Munro, E.S., Monte, M.M., Fourrier, M.C.S., Whitelaw, J., Smail, D.A., Ellis, A.E., 2010. Atlantic salmon (Salmo salar L.) serum vitellogenin neutralises infectivity of infectious pancreatic necrosis virus (IPNV). Fish Shellfish Immunol. 29, 293–297. https://doi.org/10.1016/j.fsi.2010.04.010

García-Alonso, J., Hoeger, U., Rebscher, N., 2006. Regulation of vitellogenesis in Nereis virens (Annelida: Polychaeta): Effect of estradiol-17β on eleocytes. Comp. Biochem. Physiol. A. Mol. Integr. Physiol. 143, 55–61. https://doi.org/10.1016/j.cbpa.2005.10.022

Gospocic, J., Shields, E.J., Glastad, K.M., Lin, Y., Penick, C.A., Yan, H., Mikheyev, A.S., Linksvayer, T.A., Garcia, B.A., Berger, S.L., Liebig, J., Reinberg, D., Bonasio, R., 2017. The Neuropeptide Corazonin Controls Social Behavior and Caste Identity in Ants. Cell 170, 748-759.e12. https://doi.org/10.1016/j.cell.2017.07.014

Grohmann, L., Blenau, W., Erber, J., Ebert, P.R., Strünker, T., Baumann, A., 2003. Molecular and functional characterization of an octopamine receptor from honeybee (Apis mellifera) brain. J. Neurochem. 86, 725–735. https://doi.org/10.1046/j.1471-4159.2003.01876.x

Havukainen, H., Halskau, Ø., Skjaerven, L., Smedal, B., Amdam, G.V., 2011. Deconstructing honeybee vitellogenin: novel 40 kDa fragment assigned to its N terminus. J. Exp. Biol. 214, 582–592. https://doi.org/10.1242/jeb.048314

Havukainen, H., Münch, D., Baumann, A., Zhong, S., Halskau, Ø., Krogsgaard, M., Amdam, G.V., 2013. Vitellogenin Recognizes Cell Damage through Membrane Binding and Shields Living Cells from Reactive Oxygen Species. J. Biol. Chem. 288, 28369–28381. https://doi.org/10.1074/jbc.M113.465021

Havukainen, H., Underhaug, J., Wolschin, F., Amdam, G., Halskau, Ø., 2012. A vitellogenin polyserine cleavage site: highly disordered conformation protected from proteolysis by phosphorylation. J. Exp. Biol. 215, 1837–1846. https://doi.org/10.1242/jeb.065623

Hayward, A., Takahashi, T., Bendena, W.G., Tobe, S.S., Hui, J.H.L., 2010. Comparative genomic and phylogenetic analysis of vitellogenin and other large lipid transfer proteins in metazoans. FEBS Lett. 584, 1273–1278. https://doi.org/10.1016/j.febslet.2010.02.056

Heeringen, V. J S., Veenstra, G.J.C., 2011. GimmeMotifs: a de novo motif prediction pipeline for ChIP-sequencing experiments. Bioinformatics 27, 270–271. https://doi.org/10.1093/bioinformatics/btq636

Heinz, S., Benner, C., Spann, N., Bertolino, E., Lin, Y.C., Laslo, P., Cheng, J.X., Murre, C., Singh, H., Glass, C.K., 2010. Simple Combinations of Lineage-Determining Transcription Factors Prime cis-Regulatory Elements Required for Macrophage and B Cell Identities. Mol. Cell 38, 576–589. https://doi.org/10.1016/j.molcel.2010.05.004

Huang, D.W., Sherman, B.T., Lempicki, R.A., 2009. Systematic and integrative analysis of large gene lists using DAVID bioinformatics resources. Nat. Protoc. 4, 44–57. https://doi.org/10.1038/nprot.2008.211

Hwang, S., Gou, Z., Kuznetsov, I.B., 2007. DP-Bind: a web server for sequence-based prediction of DNA-binding residues in DNA-binding proteins. Bioinformatics 23, 634–636. https://doi.org/10.1093/bioinformatics/btl672

Hystad, E.M., Salmela, H., Amdam, G.V., Münch, D., 2017. Hemocyte-mediated phagocytosis differs between honey bee (Apis mellifera) worker castes. PLOS ONE 12, e0184108. https://doi.org/10.1371/journal.pone.0184108

Jensen, P.V., Børgesen, L.W., 2000. Regional and functional differentiation in the fat body of pharaoh’s ant queens, Monomorium pharaonis (L.). Arthropod Struct. Dev. 29, 171–184. https://doi.org/10.1016/S1467-8039(00)00021-9

Klowden, M.J., Davis, E.E., Bowen, M.F., 1987. Role of the fat body in the regulation of host-seeking behaviour in the mosquito, Aedes aegypti. J. Insect Physiol. 33, 643–646. https://doi.org/10.1016/0022-1910(87)90133-8

Kucharski, R., Mitri, C., Grau, Y., Maleszka, R., 2007. Characterization of a metabotropic glutamate receptor in the honeybee (<Emphasis Type=“Italic”>Apis mellifera</Emphasis>): implications for memory formation. Invert. Neurosci. 7, 99–108. https://doi.org/10.1007/s10158-007-0045-3

Langmead, B., Salzberg, S.L., 2012. Fast gapped-read alignment with Bowtie 2. Nat. Methods 9, 357–359. https://doi.org/10.1038/nmeth.1923

Lazareva, A.A., Roman, G., Mattox, W., Hardin, P.E., Dauwalder, B., 2007. A Role for the Adult Fat Body in Drosophila Male Courtship Behavior. PLOS Genet. 3, e16. https://doi.org/10.1371/journal.pgen.0030016

Levine, B., Deretic, V., 2007. Unveiling the roles of autophagy in innate and adaptive immunity. Nat. Rev. Immunol. 7, 767–777. https://doi.org/10.1038/nri2161

Li, A., Sadasivam, M., Ding, J.L., 2003. Receptor-Ligand Interaction between Vitellogenin Receptor (VtgR) and Vitellogenin (Vtg), Implications on Low Density Lipoprotein Receptor and Apolipoprotein B/E THE FIRST THREE LIGAND-BINDING REPEATS OF VTGR INTERACT WITH THE AMINO-TERMINAL REGION OF VTG. J. Biol. Chem. 278, 2799–2806. https://doi.org/10.1074/jbc.M205067200

Li, Y., Collins, M., An, J., Geiser, R., Tegeler, T., Tsantilas, K., Garcia, K., Pirrotte, P., Bowser, R., 2016. Immunoprecipitation and mass spectrometry defines an extensive RBM45 protein– protein interaction network. Brain Res., RNA Metabolism in Neurological Disease 1647, 79–93. https://doi.org/10.1016/j.brainres.2016.02.047

Li, Z., Zhang, S., Liu, Q., 2008. Vitellogenin Functions as a Multivalent Pattern Recognition Receptor with an Opsonic Activity. PLoS ONE 3, e1940. https://doi.org/10.1371/journal.pone.0001940

Liu, R., Hu, J., 2013. DNABind: A hybrid algorithm for structure-based prediction of DNA-binding residues by combining machine learning-and template-based approaches. Proteins Struct. Funct. Bioinforma. 81, 1885–1899. https://doi.org/10.1002/prot.24330

Lycett, G.J., McLaughlin, L.A., Ranson, H., Hemingway, J., Kafatos, F.C., Loukeris, T.G., Paine, M.J.I., 2006. Anopheles gambiae P450 reductase is highly expressed in oenocytes and in vivo knockdown increases permethrin susceptibility. Insect Mol. Biol. 15, 321–327. https://doi.org/10.1111/j.1365-2583.2006.00647.x

Morishima, I., Yamano, Y., Inoue, K., Matsuo, N., 1997. Eicosanoids mediate induction of immune genes in the fat body of the silkworm, Bombyx mori. FEBS Lett. 419, 83–86. https://doi.org/10.1016/S0014-5793(97)01418-X

Münch, D., Ihle, K.E., Salmela, H., Amdam, G.V., 2015. Vitellogenin in the honey bee brain: Atypical localization of a reproductive protein that promotes longevity. Exp. Gerontol., Aging in the Wild: Insights from Free-Living and Non-Model organisms 71, 103–108. https://doi.org/10.1016/j.exger.2015.08.001

Nelson, C.M., Ihle, K.E., Fondrk, M.K., Page, R.E., Jr., Amdam, G.V., 2007. The Gene vitellogenin Has Multiple Coordinating Effects on Social Organization. PLoS Biol 5, e62. https://doi.org/10.1371/journal.pbio.0050062

Pan, M.L., Bell, W.J., Telfer, W.H., 1969. Vitellogenic Blood Protein Synthesis by Insect Fat Body. Science 165, 393–394. https://doi.org/10.1126/science.165.3891.393

Prowse, T.A., Byrne, M., 2012. Evolution of yolk protein genes in the E chinodermata. Evol. Dev. 14, 139–151.

Ramírez, F., Dündar, F., Diehl, S., Grüning, B.A., Manke, T., 2014. deepTools: a flexible platform for exploring deep-sequencing data. Nucleic Acids Res. 42, W187–W191. https://doi.org/10.1093/nar/gku365

Riesgo, A., Farrar, N., Windsor, P.J., Giribet, G., Leys, S.P., 2014. The Analysis of Eight Transcriptomes from All Poriferan Classes Reveals Surprising Genetic Complexity in Sponges. Mol. Biol. Evol. 31, 1102–1120. https://doi.org/10.1093/molbev/msu057

Roma, G.C., Bueno, O.C., Camargo-Mathias, M.I., 2010. Morpho-physiological analysis of the insect fat body: A review. Micron 41, 395–401. https://doi.org/10.1016/j.micron.2009.12.007

Salmela, H., Amdam, G.V., Freitak, D., 2015. Transfer of Immunity from Mother to Offspring Is Mediated via Egg-Yolk Protein Vitellogenin. PLoS Pathog 11, e1005015. https://doi.org/10.1371/journal.ppat.1005015

Salmela, H., Stark, T., Stucki, D., Fuchs, S., Freitak, D., Dey, A., Kent, C.F., Zayed, A., Dhaygude, K., Hokkanen, H., Sundström, L., 2016. Ancient Duplications Have Led to Functional Divergence of Vitellogenin-Like Genes Potentially Involved in Inflammation and Oxidative Stress in Honey Bees. Genome Biol. Evol. 8, 495–506. https://doi.org/10.1093/gbe/evw014

Sappington, T.W., 2002. The major yolk proteins of higher diptera are homologs of a class of minor yolk proteins in lepidoptera. J. Mol. Evol. 55, 470–475.

Seehuus, S.-C., Norberg, K., Krekling, T., Fondrk, K., Amdam, G.V., 2007. Immunogold Localization of Vitellogenin in the Ovaries, Hypopharyngeal Glands and Head Fat Bodies of Honeybee Workers, Apis Mellifera. J. Insect Sci. 7, 1–14. https://doi.org/10.1673/031.007.5201

Strand, M.R., 2008. The insect cellular immune response. Insect Sci. 15, 1–14. https://doi.org/10.1111/j.1744-7917.2008.00183.x

Tjong, H., Zhou, H.-X., 2007. DISPLAR: an accurate method for predicting DNA-binding sites on protein surfaces. Nucleic Acids Res. 35, 1465–1477. https://doi.org/10.1093/nar/gkm008

Tufail, M., Takeda, M., 2008. Molecular characteristics of insect vitellogenins. J. Insect Physiol. 54, 1447–1458. https://doi.org/10.1016/j.jinsphys.2008.08.007

Wang, R., Brattain, M.G., 2007. The maximal size of protein to diffuse through the nuclear pore is larger than 60 kDa. FEBS Lett. 581, 3164–3170. https://doi.org/10.1016/j.febslet.2007.05.082

Wheeler, M.M., Ament, S.A., Rodriguez-Zas, S.L., Robinson, G.E., 2013. Brain gene expression changes elicited by peripheral vitellogenin knockdown in the honey bee. Insect Mol. Biol. 22, 562–573. https://doi.org/10.1111/imb.12043

Xu, J., Kudron, M.M., Victorsen, A., Gao, J., Ammouri, H.N., Navarro, F.C.P., Gevirtzman, L., Waterston, R.H., White, K.P., Reinke, V., Gerstein, M., 2021. To mock or not: a comprehensive comparison of mock IP and DNA input for ChIP-seq. Nucleic Acids Res. 49, e17–e17. https://doi.org/10.1093/nar/gkaa1155

Yan, J., Kurgan, L., 2017. DRNApred, fast sequence-based method that accurately predicts and discriminates DNA-and RNA-binding residues. Nucleic Acids Res. 45, e84. https://doi.org/10.1093/nar/gkx059

