## Supplementary material for "Nuclear Translocation of Vitellogenin in the Honey Bee (*Apis mellifera*)": Figure S1 - Western blot

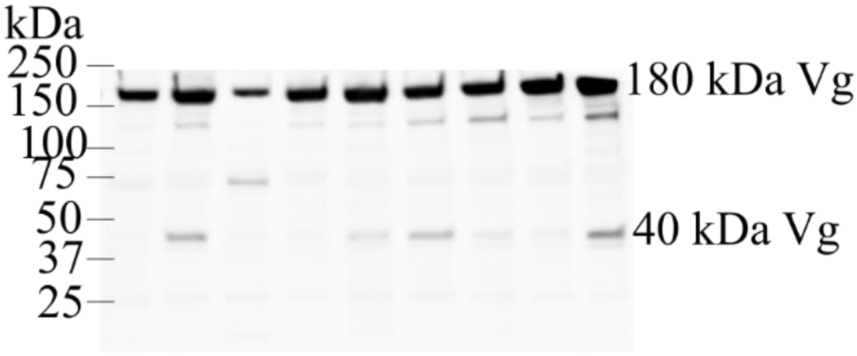


**Fig S1.** Western blot of vitellogenin N-terminal antibody (targeting amino acids 24-360) in honey bee tissue lysate samples. Abdomen of nine winter worker honey bee individuals were homogenized and western blotted (20 µg total protein per lane). There is a strong full-length vitellogenin band (180 kDa) in each sample. In addition, there are shorter fragments, whose presence and strength varies in individual samples. The previously identified 40 kDa N-terminal vitellogenin fragment is indicated. The other fragments of unknown function detected by this antibody in some individuals are ~75 kDa and ~125 kDa.
