## Supplementary Material - Additional experimental data for "Nuclear Translocation of Vitellogenin in the Honey Bee (*Apis mellifera*)"

### S2: Additional Experimental Methods and Results on Vg Cleavage Dynamics

#### Methods:

*Vg cleavage and nuclear translocation in response to immune challenge with E. coli*:

Vg is a pathogen pattern recognition receptor (Li et al., 2008, 2008; Salmela et al., 2015; Zhang et al., 2011), and since we detected Vg binding at several innate immunity genes we performed an immune challenge to explore whether bacterial exposure alters Vg cleavage and nuclear translocation patterns. First, we tested *in vitro* whether exposure to *E. coli* alters the rate of Vg cleavage. We incubated fat body protein extract rich in Vg with an increasing concentration of heat-killed *E. coli* whose bacterial proteases were inactivated by the thermal treatment (Moran et al., 2001). K-12 strain of *E. coli* were grown to $1x10^8$ cells/ml density, washed with PBS and heat-killed by shaking them in 95 °C for 30 min (Groh et al., 1996). The bacteria were then diluted 1:5 and 1:25 in PBS. 6 µl of bacteria and equal volume of 8.5 mg/ml fat body protein extract (see Salmela et al., 2015) were mixed and incubated in +26°C for 30 min. We measured the level of Vg fragmentation by Western blotting (four concentrations of *E. coli*, N = 3 each) using the Vg-N-sheet antibody (Fig S2A).

Next, we tested *in vivo* whether consumption of *E. coli* alters the rate of Vg translocation to the fat body nucleus. We used nurses bees (workers typically 4-12 days old) that were captured while tending brood, as this age/behavioral group is known to have elevated levels of Vg compared to younger or older summer workers (Amdam et al., 2005). The bees were caged in groups of 14 and fed *ad libitum* either with sterile 50 % sucrose solution or the same solution mixed with *E. coli* K-12 strain BioParticles, Alexa Fluor 488 conjugate #E13231 (Invitrogen) in a final concentration of 0.5 mg/ml. The bees were kept at 34°C in darkness for 24 h. Subsequently, the bees were immobilized by cold treatment and their fat body tissue was prepared for confocal microscopy as elsewhere in this study (see: Immunohistology). This protocol resulted in 8 successful control samples and 9 treatment samples, in addition to one antibody control sample per treatment group. The presence of the labeled *E. coli* particles in the fat body tissue was confirmed via confocal microscopy by detection of their 488 nm fluorescence signal. For each individual, we zoomed in three different fat body tissue areas with approximately 15 cells and counted the number of cells with visible nuclear localization and the number of cells without a sign of nuclear localization. Logistic regression with quasibinomial distribution was used for statistical analysis.

*Enzymatic conditions required for Vg cleavage*:

To determine how the N-sheet domain is cleaved from the Vg molecule, we searched for candidate enzymes by applying an array of protease inhibitor molecules on fat body tissue homogenate rich in Vg. Use of lysate instead of pure target protein is necessary, since pure sample has been cleared of proteases (Kondo et al., 1997). Without any protease inhibitors, full length Vg gets fragmented at room temperature in the tissue homogenate in 2 h.

Three winter worker honey bee individuals were anesthetized in cold, and their guts and ovaries were removed (see Havukainen et al., 2011). The abdomens were detached, immediately cooled in liquid nitrogen and stored in -80°C. The abdomens were homogenized in 1.5 ml ice cold PBS and insoluble material was removed by centrifugation. The inhibitors tested were (final concentration according to manufacturer's instructions): E64 (10 µM), leupeptin (100 µM), DCI (100 µM), YVAD-aomk (100 µM), EDTA (5 mM) and Roche PhosSTOP inhibitor cocktail (x 2). These are broad-range inhibitors with the exception of the highly specific caspase 1 inhibitor (YVAD-aomk), because there is a caspase 1 cut site in the Vg sequence that, hypothetically, would produce an N-terminal fragment of 40 kDa (Havukainen et al., 2011). The inhibitors were incubated with 9 µl of honey bee protein extract in PBS in total volume of 10 µl for 2 h in 28 ºC in triplicates and blotted. As controls, we had samples without inhibitors, and a sample that was kept on ice.

*Vg 3D structural model (electron microscopy)*:

Few structural models exist for Vg (Roth et al., 2013) and none of them focus on the dynamics at the N-sheet domain. To help bridge this gap, we used electron microscopy to create the most complete 3D model of insect Vg so far. Aliquots of pure Vg (1.1 mg/ml in 20 mM Tris, 150 mM NaCl, pH 7.5) were applied to carbon-coated copper grids (30 s) and stained with 2% uranyl acetate. Micrographs were taken in minimal dose conditions in a JEOL 1230 transmission electron microscope operated at 100 kV, using a low-dose protocol and a 4k x 4k TVIPS CMOS detector under the control of EM-TOOLS software (TVIPS). Final magnification of the CMOS images was 54,926. A total of 15,000 particles were selected, normalized, and CTF-corrected using procedures implemented in the XMIPP package (Scheres et al., 2008). For three-dimensional reconstructions, different starting templates were generated using the EMANstartcsym program, by common lines or using artificial noisy models and Gaussian blobs with the rough dimension of the proteins (Ludtke et al., 1999),  in both cases with identical results, which confirms the robustness of the structure obtained. The structure of the homologous protein lipovitellin from silver lamprey (1lsh) was fitted into the final vitellogenin volume and used to determine the handedness of the structure.

**Results:**

*In vitro E. coli incubation*: The presence of the bacteria appeared to enhance Vg cutting, as the amount of N-terminal fragmentation products increased in response to the number of bacteria (Fig S2A).

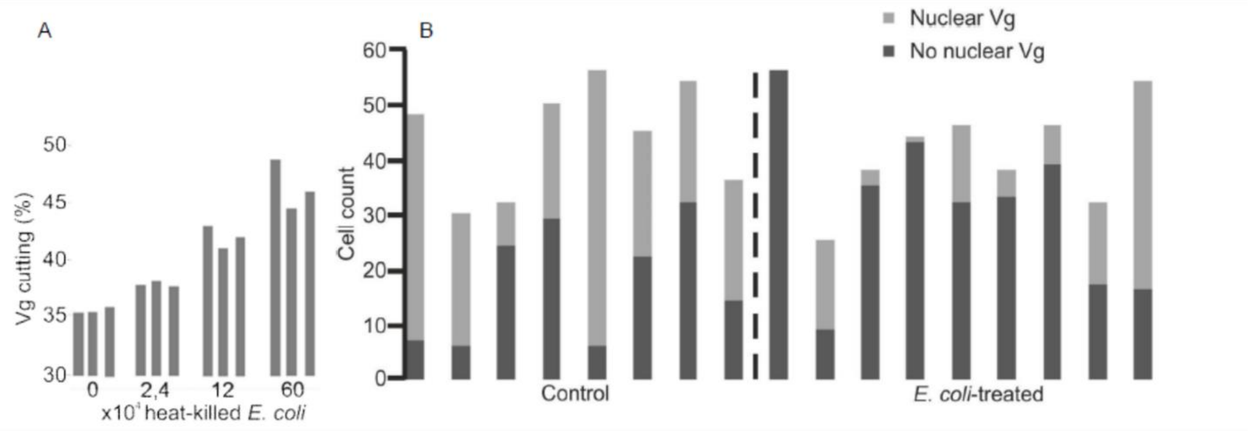


**Fig S2.** Effect of exposure to bacterial material on Vg cleavage and nuclear localization. A. Vg cleavage in response to *E. coli*. Honey bee fat body protein extract was subjected to a dilution series of heat-killed *E. coli* for 30 min in replicates of three and blotted using the Vg-N-terminal antibody. The level of Vg cutting was determined by dividing the total Vg protein signal by the fragments below 75 kDa (the nuclear fragments, see Fig 1F). B. Vg localization response to orally consumed bacteria in honey bee workers. Presence of Vg in the nucleus of fat body trophocyte cells was detected using confocal microscopy in a control group (N = 8) and in a group fed with fragments of killed *E. coli* (N = 9). Each bar represents the number of cells counted in a honey bee individual's fat body tissue.

*In vivo E. coli consumption*: Nurses fed a control diet showed nuclear translocation of Vg in 25 – 89% of their fat body trophocytes, while those fed an *E. coli* diet showed this is 0 – 64% of their fat body trophocytes (Fig S2B). Regardless of the high level of individual variation in both the control and the *E. coli*-fed group, the proportion of cells with Vg signal in the nucleus (versus cells with signal in the cytosol) was significantly higher in the control group than in the group that received bacterial fragments in their diet (b = -1.4499, z = -2.501, df = 16, p = 0.0245). These two experiments, together, indicate that the regulation of Vg cleavage and nuclear translocation responds dynamically after exposure to bacterial material.

*Enzymatic conditions required for Vg cleavage*:

Vg cutting was partially inhibited by YVAD-aomk, leupeptin (serine and cysteine protease inhibitor), EDTA (metalloprotease and phosphatase inhibitor) and phophatase inhibitor cocktail, but not by E64 (inhibitor of papain-like, but not caspase-like cysteine proteases (Rozman-Pungerčar et al., 2003)) or 3,4-dichloroisocoumarin (DCI) (serine protease inhibitor). These results show that Vg is cleaved by at least a caspase 1-like enzyme (Fig S3).

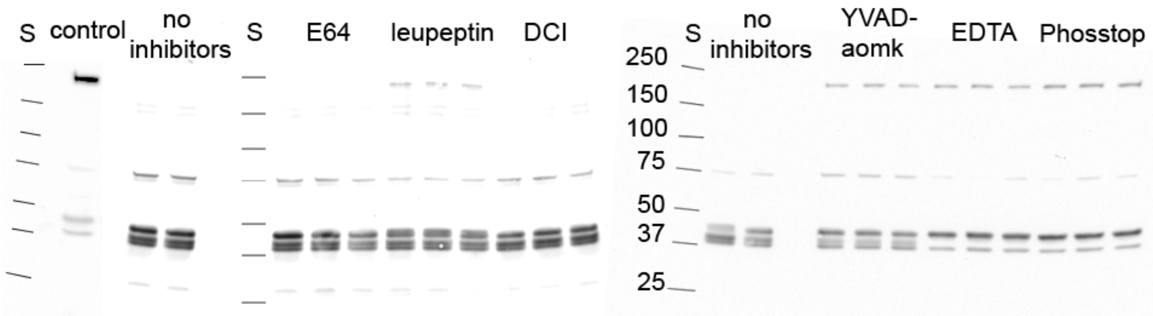


**Fig S3.** Western blot -based honey bee vitellogenin cutting inhibition assay. Un-cut, full-length vitellogenin is marked with 180 and the N-terminal domain is marked with 40 according to their size. S = size standard. The control is honey bee fat body protein extract kept on ice. The full-length vitellogenin is fully cut in 2 h in 28 °C in the absence of inhibitors (no inhibitors, N=2). The following treatments were found to prevent the cutting of vitellogenin: leupeptin, YVAD-aomk, EDTA and PhosSTOP. The treatments were done in triplicates. The figure is a combination of two blots.

*Vg cleavage site analysis via 3D structure and enzymatic assay:*

A low-resolution 3D electron microscopy reconstruction of the purified honey bee Vg was carried out using negatively stained specimens (Fig S4A), after multiple X-ray crystallization attempts failed. The structure has the general shape of a cylinder of ~140 Å high and 80 Å wide, with two small masses protruding from the top and bottom of the structure. The existing X-ray structure of the homologous protein lipovitellin from lamprey (fish) (27) was fitted into the vitellogenin volume (Fig S4B). The N-terminal domain (orange mass in Fig S4B), together with both the β-barrel and the α-helical domains of lipovitellin (yellow mass in Fig S4B), were easily fitted into the Vg volume, and the lipid-binding cavity was easily identified. The unresolved fragments of the lipovitellin structure seem to match the position of the extra masses observed in the Vg EM volume (Fig S4B). The N-sheet followed by the linker stretch appears to protrude from the structure (red mass in Fig S4B). These results suggest that enzymes have clear access to the N-sheet cleavage site. Moreover, the following treatments were found to prevent the cleavage of Vg: leupeptin, YVAD-AOMK, EDTA and PhosSTOP. This indicates that, among possibly several other molecular mechanisms, dephosphorylation (PhosSTOP inhibits dephosphorylation) and caspase-type activity (YVAD-AOMK is a specific and leupeptin a broad-range caspase inhibitor) are important for Vg cleavage to occur (Fig S3).

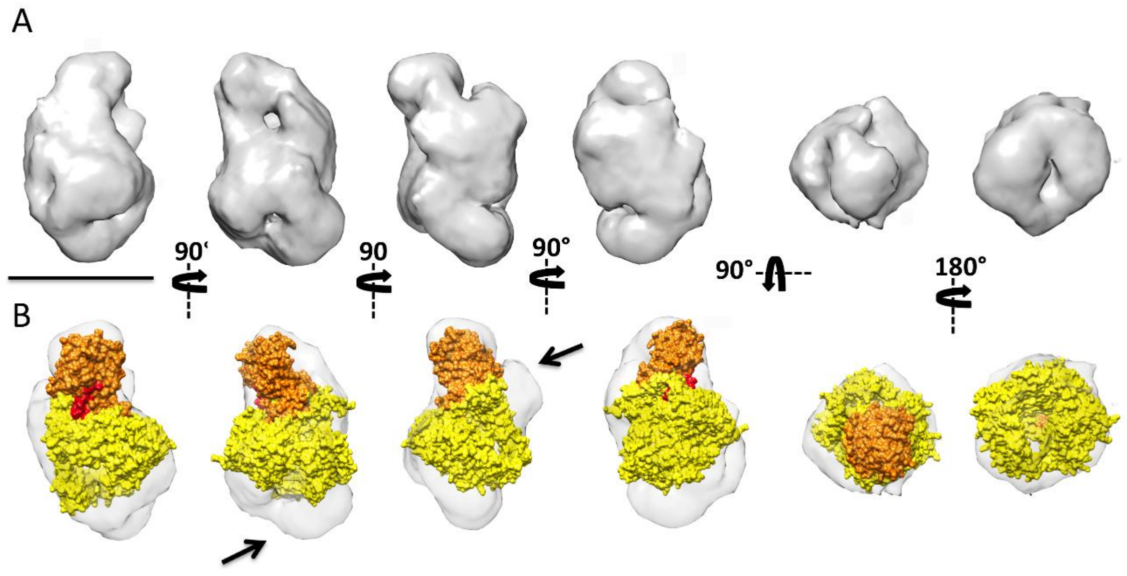

**Fig S4.** The 3D reconstruction of the honey bee Vg reveals the exposure of the cutting site. (A) Different views of the 3D reconstruction of the Vg from honey bee. The four on the left are orthogonal side views of the volume, whereas the two on the right correspond to the two end-on views of the 3D reconstruction. Bar indicates 100 Å. (B) The same views with the atomic structure of lipovitellin from lamprey (pdb 1lsh) docked into the EM 3D reconstruction. The N-sheet domain, which protrudes from the main body of the structure, is highlighted in orange. The domain colored red points to the linker that connects the N-sheet domain to the lipid cavity (yellow mass).

**Discussion:**

Given Vg's numerous immunological functions and that fact that it binds to several immune-related genes, we showed that Vg cleavage is sensitive to infection and that the system dynamically responds to immunological perturbations. Interestingly, Vg cutting is enhanced by incubation with heat-killed *E. coli*, but ingestion of *E. coli* by workers *reduces* the amount of Vg N-sheet subunit translocated to the nucleus. This finding suggests that bacterial material provokes Vg instability, but that Vg nuclear translocation is a tightly regulated process. From an immunological standpoint, it may be that Vg responds to bacterial infection by directly binding to and eliminating bacterial cells rather than up-regulating the transcription of additional immune-related genes, as Vg is known to bind to bacteria such as *E. coli* (Salmela et al., 2015) and act as a bactericidal molecule (Zhang et al., 2011). Follow-up studies are needed to fully elucidate Vg's immune response to bacterial infection, as the current study setting does not capture the acute changes caused by the bacterial challenge.

In the *in vivo* immune challenge, the level of nuclear translocation of Vg was highly variable among individuals in both the control and the *E. coli*-fed groups. Similarly, Western blots of individual bees show variation in the strength of the 40 kDa N-sheet band (S1). The hemolymph titers of full-length Vg are known to have a high level of individual variation in honey bees, which is associated with variation in stress tolerance (Seehuus et al., 2006). Similarly, variation in the nuclear Vg might be linked to certain physiological features, but this speculation needs further investigations.

This study expands our understanding of the enzymatic activity required for Vg cutting (Fig S3). Honey bee Vg N-sheet cutting site is located in a polyserine linker, whose dephosphorylation appears play a role in the proteolytic cleavage (Havukainen et al., 2012). Our inhibition assay strengthens the significance of the removal of phosphate groups, since we show that phosphatase inhibitors inhibit Vg cutting. The metalloprotease inhibitor EDTA inhibited Vg cutting similarly to phosphatase inhibitors. EDTA is known to inhibit phosphatases (Whisnant and Gilman, 2002), which may explain the result. Alternatively, EDTA might inhibit some other enzymes possibly involved in the cascade that leads into Vg cleavage. Moreover, this assay predicts that the cutting of Vg N-sheet is regulated by at least one caspase 1-like enzyme, because there is cutting inhibition by the highly specific caspase 1 inhibitor YVAD-aomk and also by leupeptin whose inhibition range covers caspases. We were led to these findings by an earlier realization that caspase 1 is the only relevant (non-gut) enzyme with a cut site in the polyserine linker (Havukainen et al., 2012). In mammals, it is well-established that caspase 1 activity is regulated by phosphorylation events (Yang et al., 2017). The mammalian caspase 1 is involved in many processes, most notably, inflammation and response to intracellular bacterial infection (reviewed by (Denes et al., 2012)). However, mammalian caspase activities do not directly translate to insects. Very little information is currently available about honey bee caspases in the literature.

Finally, we present the first 3D structure of a full-length invertebrate Vg, albeit at low resolution. The previous structural biology carried out on the Vg protein family includes an X-ray structure of lamprey lipovitellin purified from eggs (Anderson et al., 1998; Raag et al., 1988), an

NMR structure of the honey bee Vg polyserine linker (Havukainen et al., 2012), and homology modeling of honey bee Vg (Havukainen et al., 2013, 2011) and mammalian apolipoprotein B by others (Mann et al., 1999). The lamprey structure lacks four C-terminal regions because they are disordered and cannot be resolved with the X-ray diffraction (Anderson et al., 1998). Our electron microscopy structure shows the whole molecule, where the missing regions appear as protrusions surrounding the C-terminal area when compared to the lamprey X-ray structure. The electron microscopy structure reveals, for the first time, that the honey bee Vg is a monomer (see previous speculations in (Wheeler and Kawooya, 1990)), unlike most insect (Tufail and Takeda, 2008) or vertebrate Vgs (Finn, 2007). Comparison between the lamprey lipovitellin X-ray structure and our insect Vg 3D electron microscopy reconstruction reveals surprisingly little differences in the N-sheet or the linker region. The heavily phosphorylated honey bee polyserine linker is 70 amino acid residues longer than the corresponding non-phosphorylated linker in the lamprey protein (Havukainen et al., 2012). In addition, there are two insect specific loops of unknown function in the N-sheet domain (11 and 19 amino acid residues long in the *A. mellifera* Vg) (Havukainen et al., 2011). Despite the great differences in sequence and posttranslational modifications between insect and vertebrate Vgs, they appear structurally conserved. The lamprey X-ray structure docks well into the 3D reconstruction of the honey bee Vg, and shows that both N-sheet and the polyserine linker are well-exposed to solvent (Orange and red domains in Fig S4B). Such easily accessible sites are often preferred by proteases (Fontana et al., 2004), unless they are protected by some other means, for example, by phosphate groups (Cohen, 2000). Our enzyme inhibition assays suggest that dephosphorylation, indeed, may be important for Vg cleavage to occur. Furthermore, the inhibition assay suggests the Vg N-sheet domain is cleaved via caspase activity (Fig S3). However, we highlight that the cleavage and nuclear translocation of Vg might be linked to many more traits than tested or hypothesized here. We conclude that Vg cleavage might not need major conformational changes, but the phosphorylation status of Vg may play a crucial role instead.

**Works Cited:**


Amdam, G.V., Norberg, K., Omholt, S.W., Kryger, P., Lourenço, A.P., Bitondi, M.M.G., Simões, Z.L.P., 2005. Higher vitellogenin concentrations in honey bee workers may be an adaptation to life in temperate climates. Insectes Sociaux 52, 316–319. https://doi.org/10.1007/s00040-005-0812-2

Anderson, T., Levitt, D., Banaszak, L., 1998. The structural basis of lipid interactions in lipovitellin, a soluble lipoprotein. Structure 6, 895–909. https://doi.org/10.1016/S0969-2126(98)00091-4

Cohen, P., 2000. The regulation of protein function by multisite phosphorylation – a 25 year update. Trends Biochem. Sci. 25, 596–601. https://doi.org/10.1016/S0968-0004(00)01712-6

Denes, A., Lopez-Castejon, G., Brough, D., 2012. Caspase-1: is IL-1 just the tip of the *ICEberg*? Cell Death Dis. 3, e338. https://doi.org/10.1038/cddis.2012.86

Finn, R.N., 2007. Vertebrate Yolk Complexes and the Functional Implications of Phosvitins and Other Subdomains in Vitellogenins. Biol. Reprod. 76, 926–935. https://doi.org/10.1095/biolreprod.106.059766

Fontana, A., Laureto, D., Polverino, P., Spolaore, B., Frare, E., Picotti, P., Zambonin, M., 2004. Probing protein structure by limited proteolysis. Acta Biochim. Pol. 51, 299–321.

Groh, C.D., MacPherson, D.W., Groves, D.J., 1996. Effect of Heat on the Sterilization of Artificially Contaminated Water. J. Travel Med. 3, 11–13. https://doi.org/10.1111/j.1708-8305.1996.tb00689.x

Havukainen, H., Halskau, Ø., Skjaerven, L., Smedal, B., Amdam, G.V., 2011. Deconstructing honeybee vitellogenin: novel 40 kDa fragment assigned to its N terminus. J. Exp. Biol. 214, 582–592. https://doi.org/10.1242/jeb.048314

Havukainen, H., Münch, D., Baumann, A., Zhong, S., Halskau, Ø., Krogsgaard, M., Amdam, G.V., 2013. Vitellogenin Recognizes Cell Damage through Membrane Binding and Shields Living Cells from Reactive Oxygen Species. J. Biol. Chem. 288, 28369–28381. https://doi.org/10.1074/jbc.M113.465021

Havukainen, H., Underhaug, J., Wolschin, F., Amdam, G., Halskau, Ø., 2012. A vitellogenin polyserine cleavage site: highly disordered conformation protected from proteolysis by phosphorylation. J. Exp. Biol. 215, 1837–1846. https://doi.org/10.1242/jeb.065623

Kim, C.-H., Paik, D., Rus, F., Silverman, N., 2014. The caspase-8 homolog Dredd cleaves Imd and Relish but is not inhibited by p35. J. Biol. Chem. jbc.M113.544841. https://doi.org/10.1074/jbc.M113.544841

Kondo, T., Yokokura, T., Nagata, S., 1997. Activation of distinct caspase-like proteases by Fas and reaper in Drosophila cells. Proc. Natl. Acad. Sci. 94, 11951–11956. https://doi.org/10.1073/pnas.94.22.11951

Leulier, F., Rodriguez, A., Khush, R.S., Abrams, J.M., Lemaitre, B., 2000. The Drosophila caspase Dredd is required to resist Gram-negative bacterial infection. EMBO Rep. 1, 353–358. https://doi.org/10.1093/embo-reports/kvd073

Li, Z., Zhang, S., Liu, Q., 2008. Vitellogenin Functions as a Multivalent Pattern Recognition Receptor with an Opsonic Activity. PLoS ONE 3, e1940. https://doi.org/10.1371/journal.pone.0001940

Ludtke, S.J., Baldwin, P.R., Chiu, W., 1999. EMAN: Semiautomated Software for High-Resolution Single-Particle Reconstructions. J. Struct. Biol. 128, 82–97. https://doi.org/10.1006/jsbi.1999.4174

Mann, C.J., Anderson, T.A., Read, J., Chester, S.A., Harrison, G.B., Köchl, S., Ritchie, P.J., Bradbury, P., Hussain, F.S., Amey, J., Vanloo, B., Rosseneu, M., Infante, R., Hancock, J.M., Levitt, D.G., Banaszak, L.J., Scott, J., Shoulders, C.C., 1999. The structure of vitellogenin provides a molecular model for the assembly and secretion of atherogenic lipoproteins11Edited by A. R. Fersht. J. Mol. Biol. 285, 391–408. https://doi.org/10.1006/jmbi.1998.2298

Moran, A.J., Hills, M., Gunton, J., Nano, F.E., 2001. Heat-labile proteases in molecular biology applications. FEMS Microbiol. Lett. 197, 59–63. https://doi.org/10.1111/j.1574-6968.2001.tb10583.x

Raag, R., Appelt, K., Xuong, N.-H., Banaszak, L., 1988. Structure of the lamprey yolk lipid-protein complex lipovitellin-phosvitin at 2.8 Å resolution. J. Mol. Biol. 200, 553–569. https://doi.org/10.1016/0022-2836(88)90542-6

Rono, M.K., Whitten, M.M.A., Oulad-Abdelghani, M., Levashina, E.A., Marois, E., 2010. The Major Yolk Protein Vitellogenin Interferes with the Anti-Plasmodium Response in the Malaria Mosquito Anopheles gambiae. PLOS Biol. 8, e1000434. https://doi.org/10.1371/journal.pbio.1000434

Roth, Z., Weil, S., Aflalo, E.D., Manor, R., Sagi, A., Khalaila, I., 2013. Identification of Receptor-Interacting Regions of Vitellogenin within Evolutionarily Conserved β-Sheet Structures by Using a Peptide Array. ChemBioChem 14, 1116–1122. https://doi.org/10.1002/cbic.201300152

Rozman-Pungerčar, J., Kopitar-Jerala, N., Bogyo, M., Turk, D., Vasiljeva, O., Štefe, I., Vandenabeele, P., Brömme, D., Puizdar, V., Fonović, M., Trstenjak-Prebanda, M., Dolenc, I., Turk, V., Turk, B., 2003. Inhibition of papain-like cysteine proteases and legumain by caspase-specific inhibitors: when reaction mechanism is more important than specificity. Cell Death Differ. 10, 881–888. https://doi.org/10.1038/sj.cdd.4401247

Salmela, H., Amdam, G.V., Freitak, D., 2015. Transfer of Immunity from Mother to Offspring Is Mediated via Egg-Yolk Protein Vitellogenin. PLoS Pathog 11, e1005015. https://doi.org/10.1371/journal.ppat.1005015

Scheres, S.H.W., Núñez-Ramírez, R., Sorzano, C.O.S., Carazo, J.M., Marabini, R., 2008. Image processing for electron microscopy single-particle analysis using XMIPP. Nat. Protoc. 3, 977–990. https://doi.org/10.1038/nprot.2008.62

Seehuus, S.-C., Norberg, K., Gimsa, U., Krekling, T., Amdam, G.V., 2006. Reproductive protein protects functionally sterile honey bee workers from oxidative stress. Proc. Natl. Acad. Sci. U. S. A. 103, 962–967. https://doi.org/10.1073/pnas.0502681103

Tufail, M., Takeda, M., 2008. Molecular characteristics of insect vitellogenins. J. Insect Physiol. 54, 1447–1458. https://doi.org/10.1016/j.jinsphys.2008.08.007

Wheeler, D.E., Kawooya, J.K., 1990. Purification and characterization of honey bee vitellogenin. Arch. Insect Biochem. Physiol. 14, 253–267. https://doi.org/10.1002/arch.940140405

Whisnant, A.R., Gilman, S.D., 2002. Studies of reversible inhibition, irreversible inhibition, and activation of alkaline phosphatase by capillary electrophoresis. Anal. Biochem. 307, 226–234. https://doi.org/10.1016/S0003-2697(02)00062-3

Yang, J., Liu, Z., Xiao, T.S., 2017. Post-translational regulation of inflammasomes. Cell. Mol. Immunol. 14, 65–79. https://doi.org/10.1038/cmi.2016.29

Zhang, S., Wang, S., Li, H., Li, L., 2011. Vitellogenin, a multivalent sensor and an antimicrobial effector. Int. J. Biochem. Cell Biol. 43, 303–305. https://doi.org/10.1016/j.biocel.2010.11.003
